# Neuropeptides specify and reprogram division of labor in the leafcutter ant *Atta cephalotes*

**DOI:** 10.1101/2024.11.07.622473

**Authors:** Michael B. Gilbert, Karl M. Glastad, Maxxum Fioriti, Matan Sorek, Tierney Gannon, Daniel Xu, Lindsay K. Pino, Anatoly Korotkov, Ali Biashad, Josue Baeza, Richard Lauman, Anastasiia Filippova, Balint Z. Kacsoh, Roberto Bonasio, Mackenzie W. Mathis, Benjamin A. Garcia, Andrei Seluanov, Vera Gorbunova, Shelley L. Berger

## Abstract

Social insects offer powerful models to investigate the mechanistic foundation of elaborate individual behaviors comprising a cooperative community. Workers of the leafcutter ant genus *Atta* provide an extreme example of behavioral segregation among many phenotypically distinct worker types. We utilize the complex worker system of *Atta cephalotes* to test the molecular underpinnings of behavioral programming and, in particular, the extent of plasticity to reprogramming. We identify specific neuropeptides as mediators of worker division of labor in *A. cephalotes*, finding two neuropeptides associated with characteristic behaviors of leafcutting and of brood care. Manipulation via genetic knockdown or by injection of these neuropeptides led to stark loss or gain of each behavior and to transcriptomic shifts in the predicted direction, that is, towards gene pathways expressed in the natural caste. We also compare specific *A. cephalotes* worker transcriptomes with those of orthologous workers of the eusocial mammal, the naked mole rat *H. gaber*, revealing global similarities between caste-biased expression and link to specific roles of our studied neuropeptides in ants. This work underscores the essential function of neuropeptides in establishing complex social behavior and a remarkable plasticity among individual behavioral types.

## INTRODUCTION

An outstanding question in biology is how differential regulation of the genome is translated at the organismal level to regulate and alter behavior during and after development. Social insects are exceptional emerging models to investigate this question, providing a natural system exhibiting striking examples of such plasticity. Indeed, social insect colonies encompass multiple phenotypically and/or behaviorally distinct female types, despite existing as a closely related family with alternative phenotypes not arising from genetic differences between members^1^. Importantly, individual behaviors are segregated based upon phenotype^2,3^, age^4^, colony demography^5^ or a combination of these, as well as other factors, resulting in a complex reproductive division of labor with a queen and often sterile workers, or “caste” system. When polymorphism is extensive, the physical worker caste may be divided into subcastes according to allometry and trait covariance^6^. For example, worker castes perform the foraging and brood care tasks of the colony, while the reproductively active queen does not forage and primarily lays eggs. In some social insect species, exclusively among ants, tasks are further subdivided among distinct phenotypes of workers^7,8^. The molecular mechanisms that determine subcaste-specific behaviors in social insects remain understudied, despite recent findings that have begun to identify a complex system of gene and neurotransmitter regulation in activation and maintenance of these behaviors^7–14^.

One exemplar of worker division of labor is the worker subcaste system of the genus *Atta*^15,16^. The complexity of worker subcastes existing within the genus provides a robust model for investigation of mechanisms governing behavior. The intricate subcaste system in *Atta* facilitates a life history of an “agricultural lifestyle”, representing one of the most complex among ants, or, indeed, outside human society^4^. In *Atta* colonies, leaves are collected by foragers, macerated, processed, and then provided as a substrate for cultivation of a fungus that provides nutrition for the entire colony (which can reach 8 million individuals)^4^, thus establishing a multifaceted symbiotic system between the ants, two species of fungus, and a bacterium^17^. One example are workers of the species *Atta cephalotes*, which comprise multiple distinct size morphologies, among which the complex tasks of managing society are divided among workers of different sizes^16,18^. The smaller worker subcastes perform multiple distinct specialized tasks within the colony, including within-nest leaf processing^6^, management/cleaning/feeding of fungus^15^, nursing of brood^6^, and waste management^19^. In contrast, the largest subcaste morphologies regularly leave the nest, and perform the out-nest duties associated with collecting leaves^20,21^ and territorial defense^6,22^.

Neurohormones and neuropeptides are critical molecular regulators in the establishment and maintenance of social behaviors across taxa^23–25^. In humans, oxytocin is released during caretaking behaviors, such as when babies suckle mothers, to encourage milk release and overall bonding^26^. Similarly, vasopressin in humans promotes pair bonding and social recognition^26^. In the ant *Harpegnathos saltator*, which lacks worker subcastes, the neuropeptide corazonin regulates social phenotypes during transition from worker status to reproductive status, such as hunting/foraging and egg deposition^24^, an association demonstrated more recently in another ant^25^. Juvenile Hormone (JH) acts to influence social behaviors in various Hymenoptera including eusocial wasps^27,28^, bumble bees^28^, and fire ants^29^ through its gonadotropin function. The neuropeptide, Neuroparsin A (NPA) is linked to social behavior in the desert locust^30,31^. Neuroparsins have some homology to insulin peptides and may interact with the Venus Kinase Receptor (VKR)^32^; however, the precise molecular function of NPA and its role within social behavior is poorly elucidated.

Here, we uncover two novel neuropeptide/behavioral connections in *A. cephalotes* that govern principal distinct subcaste trajectories, and are able, in gain-of-function assays, to reprogram behavior. First, we identify CCAP (Crustacean cardioactive peptide), previously linked to feeding behavior in flies^48^, as central to movement of leaves, a characteristic behavior of one leafcutter worker subcaste. Second, we identify NPA, previously linked to social behavior in the desert locust, as central to the regulation of brood care, a focal behavior distinguishing the smaller subcastes from larger subcastes. We extend findings of brood care to the Naked Mole Rat (*Heterocephalus glaber*), a mammalian eusocial system, revealing striking correlations with ant forager out-nest and nurse in-nest subcastes, and strong conservation of downstream genetic pathways. We reveal NPA as a central regulator of caretaking behavior in leafcutter ants, engaging conserved pathways that are fundamental to regulation of behavior across the animal kingdom. We propose that neuropeptides serve singular functions within the intricate subcaste structure of leafcutter ants to determine complex alternative behaviors.

## RESULTS

### Behavioral and molecular characterization suggests four distinct morphological groups

We built upon prior studies of subcaste in *Atta*^15,16,33^ that utilized head width measurements, combined with behavioral observations, to distinguish behaviors associated with worker subcastes (**Fig. 1A, S1A**). While the number of distinct morphometric worker subcastes in *A. cephalotes* varies in the literature from 3^34^ to 5^18^, there is general agreement that behaviors show distinct (but partially overlapping) association with sizes: The smallest sizes perform fungal ‘gardening’^15,16,18^ and hitchhiking^35^, the smaller of the intermediate subcaste (when treated as 3 groups) performs the majority of brood care^6,18^, the larger intermediate workers perform the majority of leaf harvesting^18,20,21^, and the largest, most distinct subcaste performs defensive roles^16,18,22,36^. Further, tasks such as processing of incoming leaves^36^ and midden (trash) management^18,19^ are performed by several smaller size classes. Based upon prior work we delineated four head width ranges (**Fig. 1A, S1A**) for our own behavioral characterizations, defining four subcastes for our downstream work (**Fig. 1A**). This was done, in part, to distinguish the smaller and larger Media worker subcastes which perform distinct behavior (brood care and leaf harvesting, respectively). Animals from three genetically independent *A. cephalotes* colonies were visually observed while video recording distinct behaviors performed by each morphometric group (summarized in **Fig. 1A**). In general agreement with prior work, we found that Majors perform behaviors within and outside the nest such as defense and patrolling; Media forage for leaves through cutting and carrying behaviors; Minors perform brood care, in-nest leaf processing, and midden work (refuse management); Minims cultivate fungus hyphae and harvest hyphae, in addition to hitchhiking on leaves (thought to protect foraging Media workers from parasitoid phorid flies^35^). We chose one focal behavior from each group to evaluate for downstream experiments, and these behaviors segregated with our four morphometric groups: (1) Majors, identified by the largest gross morphology, exhibit patrolling behavior outside the nest; (2) Media execute leaf cutting and carrying; (3) Minors perform caretaking of brood; (4) Minims maintain and interact with fungal hyphae.

**Figure 1.**
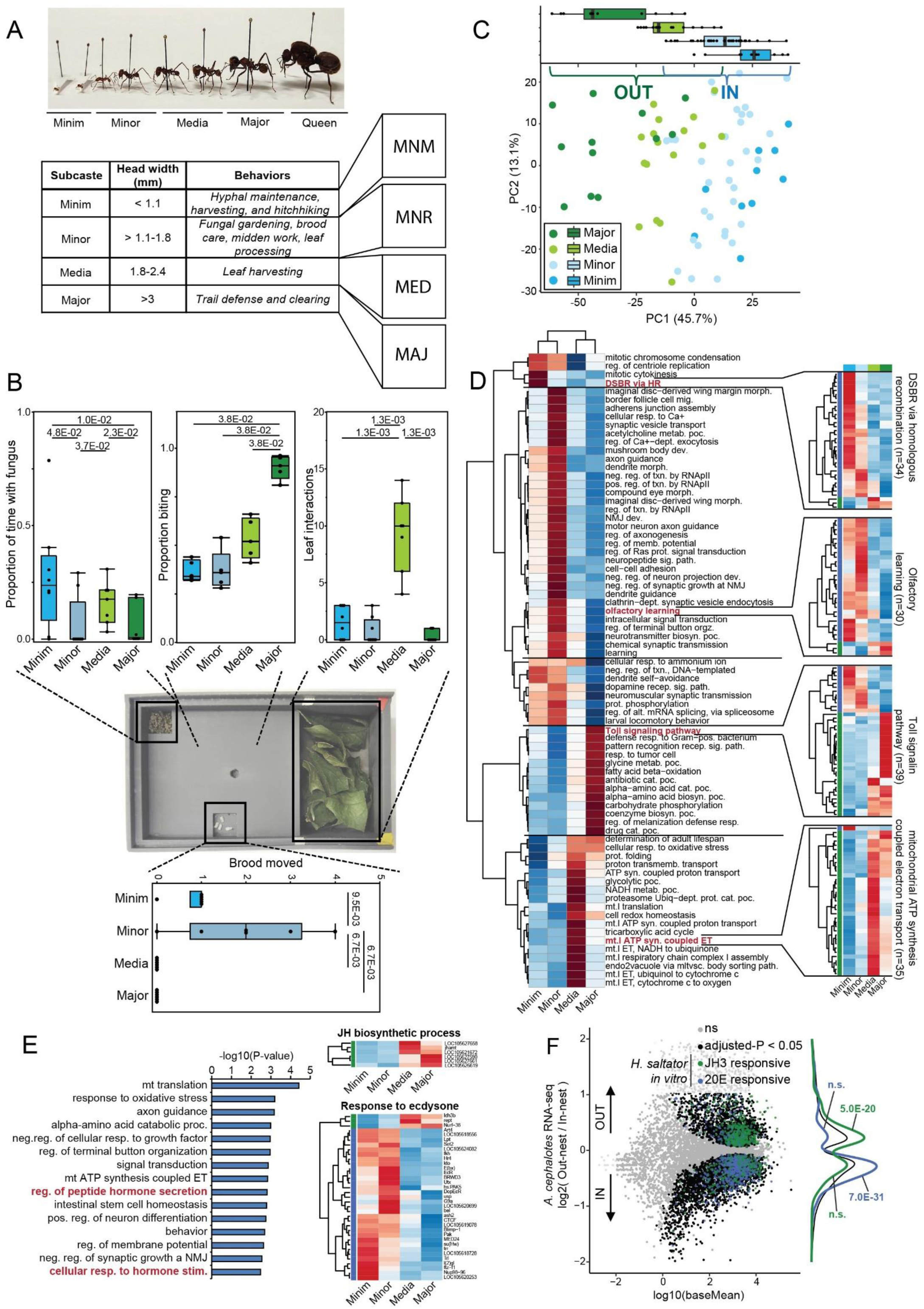
A. cephalotes shows transcriptional differentiation between major morphometric subcastes. A) Table of primary four subcaste phenotypic sizes used here, with head width range (shown in Fig. S1A), representative behaviors, and representative images of each. B) Behavior assay with baseline quantification from Automated tracking, as well as pictured representation of behavioral assay arena. Minor workers show the most interaction with fungus (upper left), while Major workers show most consistent biting response to forceps challenge (upper middle). Minor workers interact the most with brood (assessed by movements of brood to fungus; upper right; see methods) and Media workers interact the most with leaves (assessed by movements of leaves from leaf area; lower plot; see methods). C) PCA of all untreated samples, using genes showing LRT significance of ‘caste’ variable after blocking for background/prep. Upper: boxplot of PC1 values by subcaste, illustrating segregation of subcastes into major categories by this PC. D) heatmap of functional terms enriched among gene biased to a given subcaste (relative to others). Functional terms were enriched for DEGs preferentially up- or down-regulated for each subcaste (relative to others), p-values were log-transformed, then functional terms with p-values < 0.001 for enrichment among genes up-regulated in any one of the subcastes were used, and p-values representing significantly under-represented terms were given negative values for the given subcaste. Plotted to the right are representative heatmaps of gene expression (averaged among replicates) for all genes significantly differing among subcastes annotated with the given functional term. E) Functional terms enriched among genes showing significant differential expression (in either direction) when comparing “IN” subcastes (Minim+Minor) to “OUT” subcastes (Major+Media). Heatmap shows gene expression (averaged among replicate for each subcaste) for all genes significantly differing among subcastes annotated with the functional terms “Juvenile Hormone biosynthetic process” and “response to ecdysone”. F) MA plot comparing the subcastes in the two categories given by brackets (1B, upper) with JH3 and 20E LTS (*H. saltator* neuronal cultures^40^ DEGs overlapped upon these illustrating enrichment of JH3-responsive genes among ‘foraging’ subcastes and 20E-responsive genes among ‘nest’ subcastes. p-values labeling distributions represent significance of overlap between a given hormone’s DEGs and genes biased to the given group from a fisher’s exact test comparing groups.

Based upon our preliminary observations of baseline behaviors for these key subcastes we established standardized assays using single ants to capture and quantify primary distinguishing behavioral elements within the colony. We utilized an arena with three sections with separate contents to measure interactions within each: fungus from the parent colony, pupae as brood, and leaves (**Fig. 1B**, center panel). Ants were introduced into the arena and videos were recorded for 20 minutes (following a five-minute acclimation period containing all contents). We performed automated tracking of each subcaste within the behavioral assays, and quantified interactions and baseline locomotion (**Fig. 1B**). Evaluation of these four morphometric subcastes using this assay validated its utility: (1) Majors and Media never interacted with brood, whereas Minors and Minims picked up and moved pupae to the fungus (**Fig. 1B**, bottom panel). (2) Media consistently interacted with leaves whereas Majors did not (**Fig. 1B**, top right). (3) Minims exhibited the longest interaction time with fungus (**Fig. 1B**, upper left). (4) Majors showed highest frequency aggression via a blinded biting assay (**Fig. 1B**, top center) – manually scored due to their limited engagement with the other behavioral components in the assay. Thus, our assays captured several subcaste-related key behavioral characteristics of leafcutter ants (for examples, see **Supplemental videos S1-S4**).

We initiated molecular characterization of the subcastes via RNAseq to obtain whole brain transcriptomes of Major, Media, Minor, and Minim individuals, as defined above (**Fig S1A**), collected while performing distinct behaviors within the colony. For this initial characterization we extensively sampled to include additional distinct colony behaviors, resulting in eight sample types: gardening (Minims), hitchhiking, in-nest leaf processing, midden work, brood care (Minors), carrying and cutting of leaves (Media), and patrolling (Majors). We found only slight transcriptomic differentiation after splitting by these eight behaviors, with few DEGs specific to a given behavioral group (**Fig. S1B; Fig. S1C**, upper), however, we found that the transcriptome was far more informed by morphometric subcaste than behavioral group (see **Fig. 1A**), with this approach revealing many differentially expressed genes (DEGs) biased to each subcaste (**Fig. S1C**, lower). Indeed, PCA analysis of samples using overall subcaste-differing genes (likelihood ratio test of genes differing between any subcaste; adjusted P < 1E-04; n=2,111) revealed clear partitioning of samples by morphometric groups (**Fig. 1C**). This showed demarcation of the samples associated with our morphometric grouping, generally delineated by Major (defense and patrolling), Media (foraging), Minor (brood care and nest tasks), or Minim (fungal maintenance) (**Fig. 1C**; note PC1, top). These behaviors could then be separated most broadly in binary fashion as “in-nest” (Minim and Minor, subcastes that do not typically leave the nest) vs “out-nest” (Media and Major, subcastes that exit the nest) behaviors (**Fig. 1C**, IN vs OUT). While hitchhiking Minims do exit the nest, these were included due to the lack of transcriptomic differences from the other Minim group (**Fig S1C**) and stationary behavior during hitchhiking. Identification of genes upregulated in each subcaste relative to the others showed that each morphometric subcaste possessed many uniquely elevated genes (>1,000 genes per subcaste; GO categories in **Fig. 1D**, left; example genes in **Fig. 1D**, right, full list in **Table S1**). We noted certain distinct functional terms enriched for DEGs within each worker subcaste (**Fig. 1D**, left to right): Minims were enriched for DNA damage response and neuronal function; Minors were enriched for numerous terms of neuronal function and cognition; Media were enriched for multiple metabolic terms related to mitochondrial function and glycolysis; Majors were enriched for catabolic processes and innate immune signaling (**Fig. 1D**). In addition, we found strong subcaste expression bias of 297 genes annotated with a DNA binding domain (see Methods), further supporting distinct transcriptional states of subcastes apart from the “in-nest”/”out-nest” separation (**Table S2**, top ten transcription factors per subcaste in **Fig. S1D**). For example, SoxN and Lmx1a were biased to Minors consistent with functional terms related to neuronal function^37^, whereas Majors showed highest expression of sugarbabe (sug;^38^) and Mitf^39^, consistent with catabolic terms associated with Major-biased genes (**Fig. S1D**). These results suggest that the four distinct morphological subcastes that differ in characteristic behavior, also differ in brain gene expression.

We noted, in particular, that “response to hormone stimulus” and “hormone secretion” appeared in the top 15 functional terms (**Fig. 1E**, left) enriched among genes differing between any subcaste (as **Fig. 1D**). We thus analyzed hormone regulation in the subcastes, starting with Ecdysone (20E) and Juvenile hormone (JH3), because prior work by our group and others identified these as instrumental in distinguishing reproductive/non-forager compared to worker/forager, respectively^7,8,40^. Consistently, of subcaste DEGs, genes annotated with “response to ecdysone” (32/35; 91%) were biased to in-nest subcastes (Minim and Minor) (**Fig. 1E**, lower right), whereas all 6 DEGs assigned “Juvenile hormone biosynthetic process” were biased to out-nest subcastes (Media and Major) (**Fig. 1E**, upper right) (see **Table S1**). To both further assess this—and to evaluate whether this pattern globally compares to patterns found between castes of other ants where the hormones JH3 or 20E are associated with division of labor^9,40^—we compared DEGs between out-nest and in-nest subcastes in *A. cephalotes* to DEGs from our previous *in vitro* stimulation of *H. saltator* neuronal cultures with JH3 or 20E^40^. We observed strong overlap with JH3-upregulated genes in *H. saltator* cultured cells enriched among out-nest workers (Major/Media) in *A. cephalotes*, and, in contrast, 20E-upregulated genes in *H. saltator* cultured cells were enriched among in-nest workers in *A. cephalotes* (**Fig. 1F**). Further comparison of DEGs biased within the four major morphometric *A. cephalotes* subcastes to DEGs of hormonal stimulation in *H. saltator* revealed Media workers most strongly correlating with JH3-induced genes (**Fig. S1E,** upper, light green) and Minor most strongly associating with 20E-biased genes (**Fig. S1E**, upper middle, light blue). We also compared *A. cephalotes* to our previous RNA-seq data from Florida carpenter ant *C. floridanus* Major and Minor worker brain at eclosion (day 0; d0) and after five days (d5)^7^, to discern how the more complex four morphometric subcaste system in *A. cephalotes* compared to a simpler system with only two worker subcastes. We found that genes biased to Media and Major workers in *A. cephalotes* showed elevated expression in *C. floridanus* d0 Major workers (**Fig. S1E**, lower middle, light and dark green), despite Minor workers being responsible for foraging in *C. floridanus*^7,9,41^; however when comparing to d5 Major and Minor *C. floridanus* data the expected association was observed, with genes biased to *A. cephalotes* Media workers showing the strongest bias to *C. floridanus* Minor workers (**Fig. S1E**, lower, light blue). Overall, we found logical similarities between the more morphologically elaborated subcastes of *Atta* with the less morphologically and behaviorally elaborated subcaste systems of *H. saltator* and *C. floridanus*. More specifically, we found that specific *A. cephalotes* worker subcastes correlate best with *C. floridanus* (cf) functional subcastes (Minor to cfMinor, Media to cfMajor), consistent with associations with hormonal signaling established in both *H. saltator*^40^ and *C. floridanus*^9^. This supports conservation of association between hormonal signaling and caste across ants^40,42^, as well as partial consistency with worker subcaste in *C. floridanus*, which also shows consistent association with hormonal signaling^9^. Nevertheless, these associations are insufficient to explain segregation into >2 morphological and behaviorally distinct worker types found in *A. cephalotes*.

### Neuropeptides distinguish the four morphological groups

Thus, we more deeply investigated neuropeptides towards elucidating the basis of the more complex behavior among the four morphological subcastes of *A. cephalotes*, beyond the strong binary association of Major+Media with JH3 regulation, and Minor+Minim with 20E regulation. Given that functional terms for peptide hormone secretion were among the top subcaste DEGs (see **Fig. 1D**), we examined gene homologues of insect neuropeptides or neuropeptide receptors differing between in-nest (Minim+Minor) and out-nest (Major+Media) subcastes and found many neuropeptides and receptors exhibiting differential expression biased to these two main in-nest and out-nest subcaste groups (indicated in **Fig. 2A**). Further, inspection of differential expression among the four morphometric subcastes identified a large number of neuropeptides or neuropeptide receptors showing significant bias to at least one subcaste, with multiple neuropeptides most highly expressed in one of the subcastes (**Fig. 2B**, upper panels). For example, we identified *sNPF*, a neuropeptide that regulates food intake and body size in *Drosophila*^43^, as specifically upregulated in Minim (**Fig. 2B**, blue; full set of peptide hormones and receptors in **Fig. S2A**), which may relate to Minim interaction with the colony fungus food source (see **Fig. 1B**). *Dh31*, a neuropeptide that regulates body temperature preference and sleep in *Drosophila*^44^, was most highly expressed in Minor (**Fig. 2B**, light blue), which may connect to Minor exclusive in-nest behavior. The leaf management Media subcaste showed the highest expression of *CCAP*, *Capa*, *SIFa*, and Partner of Bursicon (*Pburs*)^45^ (**Fig. 2B**, light green); many of these neuropeptides are implicated in regulating feeding behavior: *Capa* in regulating energy balance and feeding^46^, *SIFa* in circadian feeding rhythms^47^, and CCAP in regulating feeding behavior and circadian rhythm^48,49^. Lastly, among neuropeptides elevated in Major, Neuroparsin-A (*NPA*), a neuropeptide originally characterized in regulation of gregarious behavior in the Desert Locust and lost in *Drosophila melanogaster*^30,50^, exhibited striking bias to Major workers along with *ilp1* and *CNMa*^51^ (**Fig. 2B**, dark green). We also detected multiple neuropeptide receptors with similar subcaste-bias, notably the putative receptor for *NPA*—homolog to *VKR*^32^— showing strong bias to Major (**Fig. 2B**, lower panels). In addition to these specific subcaste-biased neuropeptides, PCA analysis focusing on neuropeptide expression (**Fig. S2B**) showed partial recapitulation of main subcaste in-nest and out-nest groups (**Fig. S2B**, note PC2, right).

**Figure 2.**
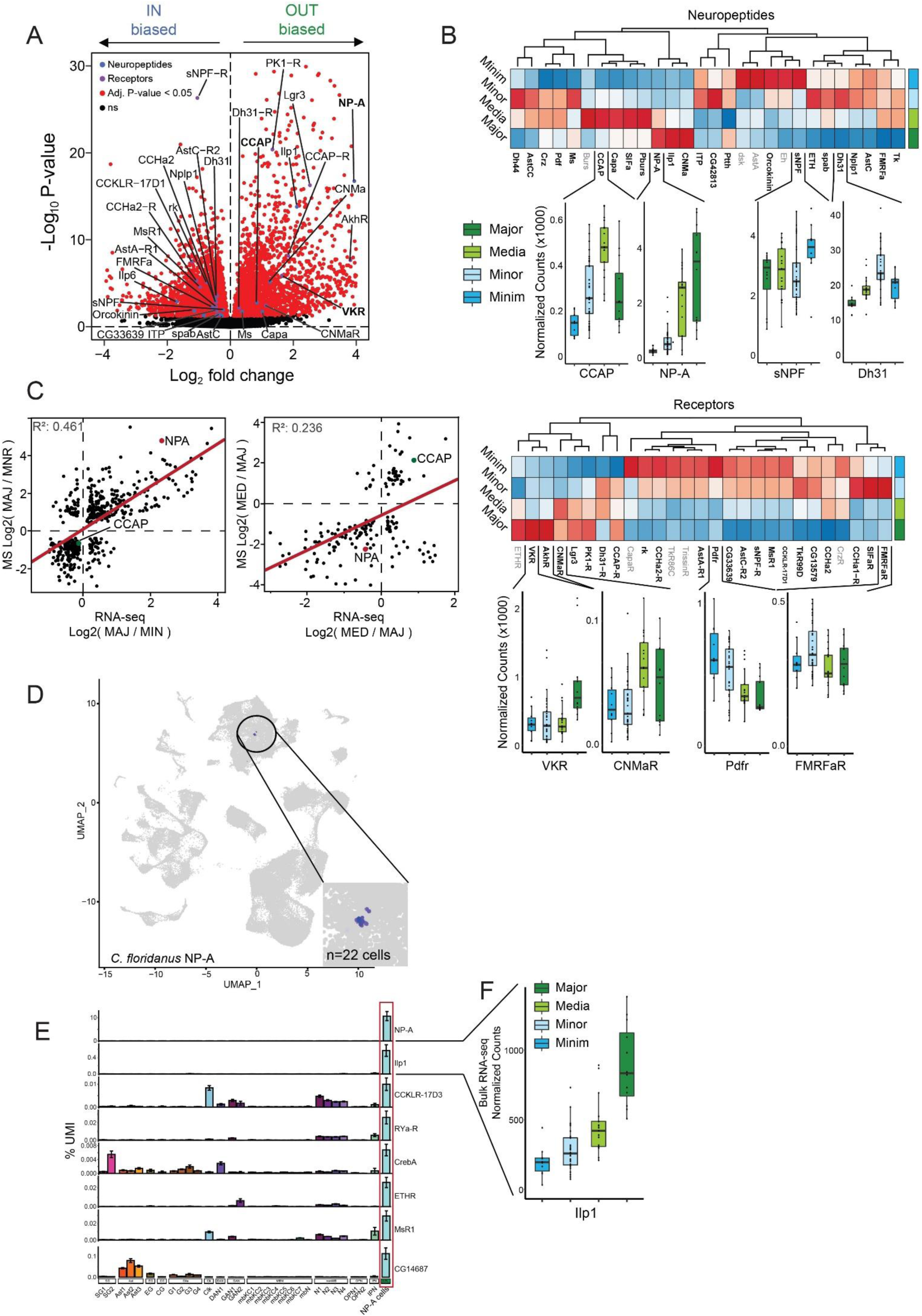
Neuropeptide expression distinguishes morphometric worker castes in A. cephalotes. A) Volcano plot showing differentially expressed genes between out-nest and in-nest subcastes. Genes with homology to know insect neuropeptides or receptors are colored in blue or purple dots, respectively. B) Heatmaps of replicate-averaged expression for each of the main *A. cephalotes* subcastes here for putative (top) neuropeptides and (bottom) neuropeptide receptors. Gene names in bold represent genes significantly differing among subcastes. For each subcaste plots of normalized counts for a representative neuropeptide and receptor showing significant highest expression in the given subcaste is given below the respective heatmaps. C) Scatterplot of RNA-seq expression difference (log2 fold change) between Major vs Minor (left) and Media vs Major (right) compared to the log2 ratio of whole proteome mass-spectrometry comparing the same subcastes. NPA is highlighted in red showing high levels in Major (as compared to Minor) in both MS and RNA-seq. Genes plotted are limited to those showing consistent significant difference between the compared subcastes in both RNA-seq and MS. D) UMAP plot from *C. floridanus* illustrating cells expressing NPA (minimum expression > 4; 22 cells). E) Genes showing highest expression in NPA cells as compared to all other clusters in scRNA-seq from *C. floridanus*^9^, illustrating NPA cells show high levels of multiple neuropeptides or receptors. F) Ilp1 exhibits highest expression in Major morphometric group by bulk RNA-seq.

To better elucidate and solidify the expression patterns of neuropeptides among *A. cephalotes* subcastes, we performed global proteome mass spectrometry from brains of all worker types. The correlation between the proteome and transcriptome of two focal groups (Major/Minor; out-nest/in-nest) showed *NPA* prominently biased to Major compared to Minor at both the transcription and protein levels (**Fig. 2C**, left; **Fig. S2C**; see **Table S3**); in contrast, we noted that *CCAP* was strongly biased to Media (leaf management) relative to Majors (**Fig. 2C**, right). We more broadly contextualized the bias in expression of *NPA* to out-nest subcastes by evaluating expression across several Hymenopteran species. We compared forager to nurse (*M. pharaonis*, *S. invicta*, *B. terrestris*), or Major to Minor (*C. floridanus*), and most of these comparisons (75%) showed elevated NPA in forager/Major workers (**Fig. S2D**). Because NPA was elevated in Majors of *C. floridanus* similarly to *A. cephalotes* Majors, we evaluated single-cell RNA sequencing data from *C. floridanus* brain (**Fig. S2E**)^9^. We found *NPA* exclusively expressed in few (n=22) cells (**Fig. 2D**), which—given other known genes expressed in the region^52^—likely correspond to the pars intercerebralis, a brain neurosecretory region critical for neuropeptide production. Within *C. floridanus* cells expressing *NPA*, several other neuropeptides or neuropeptide receptors also showed highest expression, including Ilp1, a peptide hormone also most highly expressed in Majors, suggesting possible crosstalk between *NPA* and insulin signaling (**Fig. 2E**, left and right), which is supported by co-expression of the putative *NPA* receptor (*VKR*) and the insulin receptor (*InR*) in several glial types including perineurial glia (**Fig. S2F**). Taken together, transcriptomic meta-analyses of other ants with subcaste-based division of labor supports the importance of individual neuropeptides to behavior, and in particular, *NPA*.

Similarly, *CCAP*, the neuropeptide showing the most significant bias to Media, was expressed in *C. floridanus* scRNA-seq data within even fewer cells than NPA (n=16)^9^, and these cells showed a strong signal of expression of *Dh31* and other genes regulating circadian rhythm (*cyc*, *cwo*, *Pdfr*, *per*, *Pdp1*; **Fig. S2G** and **S2H**). This association suggests crosstalk between *CCAP* and circadian regulation, consistent with the highly circadian nature of *A. cephalotes* leaf foraging behavior^53^. Together, our results suggest that specific neuropeptides, in the context of the general JH3/20E delineation of primary subcaste groups (**Fig. 1F**), may facilitate the advanced behavioral differentiation among *A. cephalotes* subcastes.

### NPA and CCAP control distinct subcaste-specific behaviors

We probed behaviors associated with subcaste-specific expression of neuropeptides in *A. cephalotes*, utilizing the behavioral assays above to record each ant once (see **Fig. 1B**), coupled with genetic and neuropeptide perturbations. Given the morphological and neuropeptide disparities between the Major and Minor groups, we investigated whether neuropeptide expression is connected to their focal behaviors, concentrating on neuropeptides that were most strikingly different at both the transcriptomic and proteomic levels. The top two neuropeptides we chose to investigate, NPA and CCAP, showed the highest correlation in RNA compared to protein expression levels, in addition to being the most differentially expressed neuropeptides at the protein level (see Figs 2C and S2C). We hypothesized that Neuroparsin-A (NPA) elevation in the out-nest Major and Media subcastes (**Fig. 2B**) might exert suppression of brood care, which is a primary behavior of the in-nest Minor and Minim subcastes, and notably both exhibited low NPA levels. Indeed, Major and Media spend little time with brood (see **Fig. 1B**, bottom) and only Minor (highest level) and Minim move brood to fungus (**Fig. 3A**), an important behavior related to brood care^10^. Indeed, these behaviors are central to forager/nurse delineation broadly in insects in addition to other species (e.g. crustaceans) where neuropeptide expression is linked to specific behaviors. Thus, we investigated the importance of both NPA and its putative receptor VKR to suppression of caretaking. We initially tested the Major subcaste, because it exhibited the highest NPA expression (see **Fig. 2B**) and strongest transcriptomic distinction from other subcastes (see **Fig. 1B**). We injected dsiRNA (double-stranded inhibitory RNA) targeting *NPA*, or targeting its putative receptor *VKR*, into Major brain, validated RNA knockdown (**Fig. S3A**), and performed the behavioral assay 24 hours post-injection, at a timepoint of strong reduction of NPA (**Fig. S3A**). We found that knockdown of either *NPA* (red, n = 10, p-value = 5.2E-03) or *VKR* (orange, n = 10, p-value = 3.5E-02) led to significant increase in Major picking up the brood in their mandibles and depositing brood onto the fungus (**Fig. 3B**; **Supplemental video S5**). Of particular note was that NPA knockdown in Major led to similar levels of brood movement to fungus compared to natural baseline levels of brood movement detected in Minor (compare **Fig. 3B** Major, red to **Fig. 3A** Minor, light blue). Thus, in Major ants, the key defensive subcaste, reduction of NPA or VKR led to substantial acquisition of caretaking.

**Figure 3.**
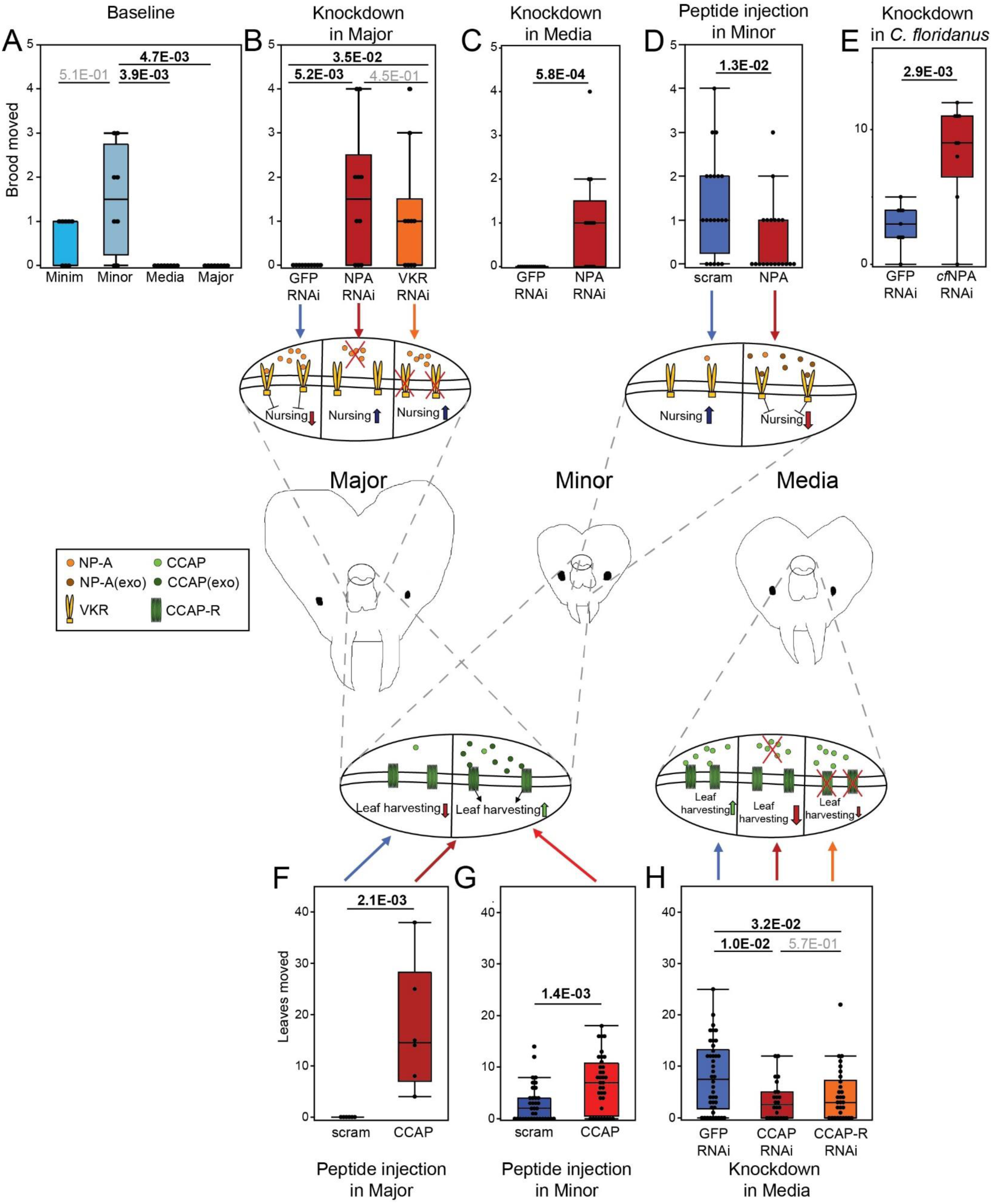
Behavioral changes with NPA and CCAP. A) Baseline levels of brood movement behavior among untreated castes. P-value generated by a Wilcoxon signed-rank test with Bonferroni correction following a significant Kruskal-Wallis test across all groups (n=8, each subcaste). B) Brood movement behavior following NPA and VKR KD in Major workers. P-value generated by a Wilcoxon signed-rank test with Bonferroni correction following a significant Kruskal-Wallis test across all groups (n=10, each sample type). C) Brood movement behavior following NPA KD (or control GFP KD) in Media workers. P-values generated a Mann-Whitney U test (n=15, each sample type). D) Brood movement behavior following NPA or control (scrambled peptide) injection in Minor workers. P-values generated a Mann-Whitney U test (n=20, each sample type). E) Brood movement behavior following NPA KD (or control GFP KD) in *C. floridanus* Major workers. P-values generated a Mann-Whitney U test (n=9, each sample type). F) Leaf movement behavior following CCAP or control (scrambled peptide) injection into Major workers reflecting stark acquisition of leaf movement behavior with CCAP injection. P-values generated a Mann-Whitney U test (n=6, each sample type). G) Leaf movement behavior following CCAP injection into Minor, showing ∼acquisition of leaf movement in alternative subcaste upon CCAP injection. P-values generated a Mann-Whitney U test (n=36, each sample type). H) Leaf movement behavior following CCAP and CCAP-R KD. P-value generated by a Wilcoxon signed-rank test with Bonferroni correction following a significant Kruskal-Wallis test across all groups (GFP n=42, CCAPi n=24, CCAP-Ri n=30).

Using proteomic approaches we further investigated whether VKR is a *bona fide* receptor for NPA. We analyzed the protein interactome of NPA. Brain lysate from Majors was incubated with FLAG-tagged purified recombinant NPA or FLAG-tagged scrambled peptide as a negative control (**Fig. S3B**), then immunoprecipitated with FLAG antibody, followed by mass spectrometry using spike-in of a heavy-labeled VKR peptide to assist in identification of VKR. VKR was a top interacting protein and, notably, was the only interacting receptor (**Fig. S3C**; all interacting proteins in **Table S4**). These findings further support VKR as a receptor for NPA and indicate that the similar behavioral reprogramming of Major to caretaking using dsiRNA to reduce either NPA or VKR described above was due to their molecular pathway.

We further tested the role of NPA in caretaking behavior, probing whether it also suppressed caretaking in Media, which, similar to Majors, have high levels of NPA (**Fig. 2B** upper), interact far less with brood than Minim (**Fig. 1B** lower), and do not move brood (**Fig. 3A**). Again, dsiRNA reduction of *NPA* (**Fig. S3A**) led to increased brood movement by Media (**Fig. 3C**, n = 13, p-value = 5.8E-04), thus indicating a general role of NPA in governing caretaking. As a further important test of the proposed suppressive role of NPA peptide in caretaking, we used recombinant *A. cephalotes* NPA (**Fig. S3B**) to perform brain injection into Minor, which exhibited the most robust brood interaction and movement (see **Fig. 1B**). NPA injection led to strong reduction of brood movement by Minors (**Fig. 3D**, n = 20, p-value = 1.8E-02), and, importantly, there was no difference in mortality between injection of scrambled control peptide and NPA (**Fig. S3D**). As discussed above, NPA levels were also elevated in Majors of *C. floridanus* (see **Fig. S2D**), hence we tested conservation of NPA in regulating caretaking via dsiRNA (validation in **Fig. S3E**). We observed increased brood movement with *NPA* knockdown (**Fig. 3E**, n = 9, p-value = 2.9E-03), suggesting a conserved role of NPA in suppressing caretaking across ants.

We investigated—in addition to suppression of caretaking behavior—whether NPA can also promote the acquisition of a behavior. We examined biting, an aggressive behavior associated with natural Majors (**Fig 1B**) and found that Major biting following *NPA* KD was significantly decreased (n=, p=1.3E-03; **Fig S3F**, left). Conversely, Minor worker biting was increased when injected with recombinant NPA, supporting that NPA promotes aggression as in natural Majors (n=, p=9.1E-03; **Fig S3F**, right). In addition, we evaluated the impact of social context on the effects of NPA, via *NPA* KD in Majors. We performed the behavioral assay as before, but included an equivalent number of Minor workers in the assay, either in air-permeable cages or free to roam the arena as Majors, and found no alteration of the increase in caretaking behaviors in Major workers via *NPA* KD (**Fig S3G**). Thus, reduction of NPA in Majors promotes caretaking regardless of presence of the typical caretaking caste (Minors).

A remarkable feature of the leafcutter ant species is their elaborate subcaste-specific behaviors. We investigated CCAP as a second neuropeptide that may specify the eponymous distinct behavior of leaf management. *CCAP* is elevated in Media (see **Fig. 2B**) and is implicated in *D. melanogaster* feeding behavior^48^. We tested CCAP in governing harvesting and transport of leaves, via injection of recombinant CCAP peptide into Major and Minor, which do not exhibit leaf collection (see **Fig. 1B**). We evaluated movement of leaves over 24 hours following a one hour acclimation period in the arena containing all contents. CCAP injection resulted in acquisition of leaf movement (as defined by an ant picking up a leaf fragment in its mandibles and placing the leaf elsewhere in the behavior chamber; see Methods) by both Major (**Fig. 3F**, n = 6, p-value = 2.1E-03, **Supplemental video S6**) and Minor (**Fig. 3G**, n = 36, 1.4E-03). Given the distinct transcriptomes and baseline behaviors of Major and Minor subcastes (see **Fig. 1B, 1C**), this similar acquisition of leaf movement strongly supports the hypothesis that CCAP is a key regulator of leaf harvesting behavior. Conversely, *CCAP* knockdown in Media (**Fig. S3H**, middle) resulted in significant decrease in leaf movement (**Fig. 3H**, red, n = 24, p-value = 1E-02), and knockdown of CCAP receptor (*CCAP-R*) (**Fig. S3H**, right) resulted in a trend towards decreased leaf movement (**Fig. 3H**, orange, n = 30, p-value = 3.2E-02).

We validated the behavioral acquisition perturbations for both NPA (KD) and CCAP (peptide injection) using automated tracking of control- or treatment-injected ants. Automated video tracking supported our results, with *NPA* KD in Majors resulting in significant increase in Major workers at brood and at fungus but did not significantly alter leaf interactions or locomotion (**Fig S3I**, left). This shows *NPA* KD results in generally higher residence time at/around both brood and fungus (we note that the automated tracking does not allow detection of discrete transport of brood to fungus). Similarly, automating tracking showed that peptide injection of CCAP resulted in significant increase in leaf carrying, but no significant change in interaction with fungus or locomotion (**Fig S3I**, right). Further, across 2h of video tracking throughout the assay, Majors injected with CCAP continued to move leaves (**Fig S3J**, left), and spent more time in the area containing the leaves compared to control-injected Majors (**Fig S3J**, right).

Taken together, our results reveal that neuropeptides facilitate and can reprogram elaborate subcaste-specific behavior in the complex social system of *A. cephalotes*. We find that NPA negatively regulates caretaking behavior in out-nest subcastes Major and Media, whereas CCAP positively regulates leaf harvesting behavior in in-nest subcastes Minor and Minim.

### NPA and CCAP reprogram the worker transcriptome, paralleling behavioral changes

We explored gene regulation underpinning the behavioral specification associated with NPA and CCAP reprogramming, via transcriptomic analyses of the perturbations described above. We analyzed NPA in Major via brain dsiRNA knockdown of *NPA* and of its putative receptor *VKR*, followed by RNA-seq of brains 6-, 12-, and 24-hours post-knockdown (validation in **Fig. S3A**). Similar overall transcriptomic outcomes resulted from either knockdown, with similar genes up-regulated or down-regulated following *NPA* KD compared to *VKR* KD (**Fig. S4A**, positive correlations of left three columns). We note that these results further support VKR as the receptor for NPA, as genes up- and down-regulated following *NPA* KD generally showed the same response following *VKR* KD (**Fig S4A**). PCA analyses of *NPA* and *VKR* knockdown in Major, compared to knockdown of *GFP* as a negative control, showed marked shifting of the Major transcriptome towards natural Nurse along PC1 (**Fig. 4A**, x-axis; VKR **Fig. S4B;** grey shifting to red, similar to blue on PC1).

**Figure 4.**
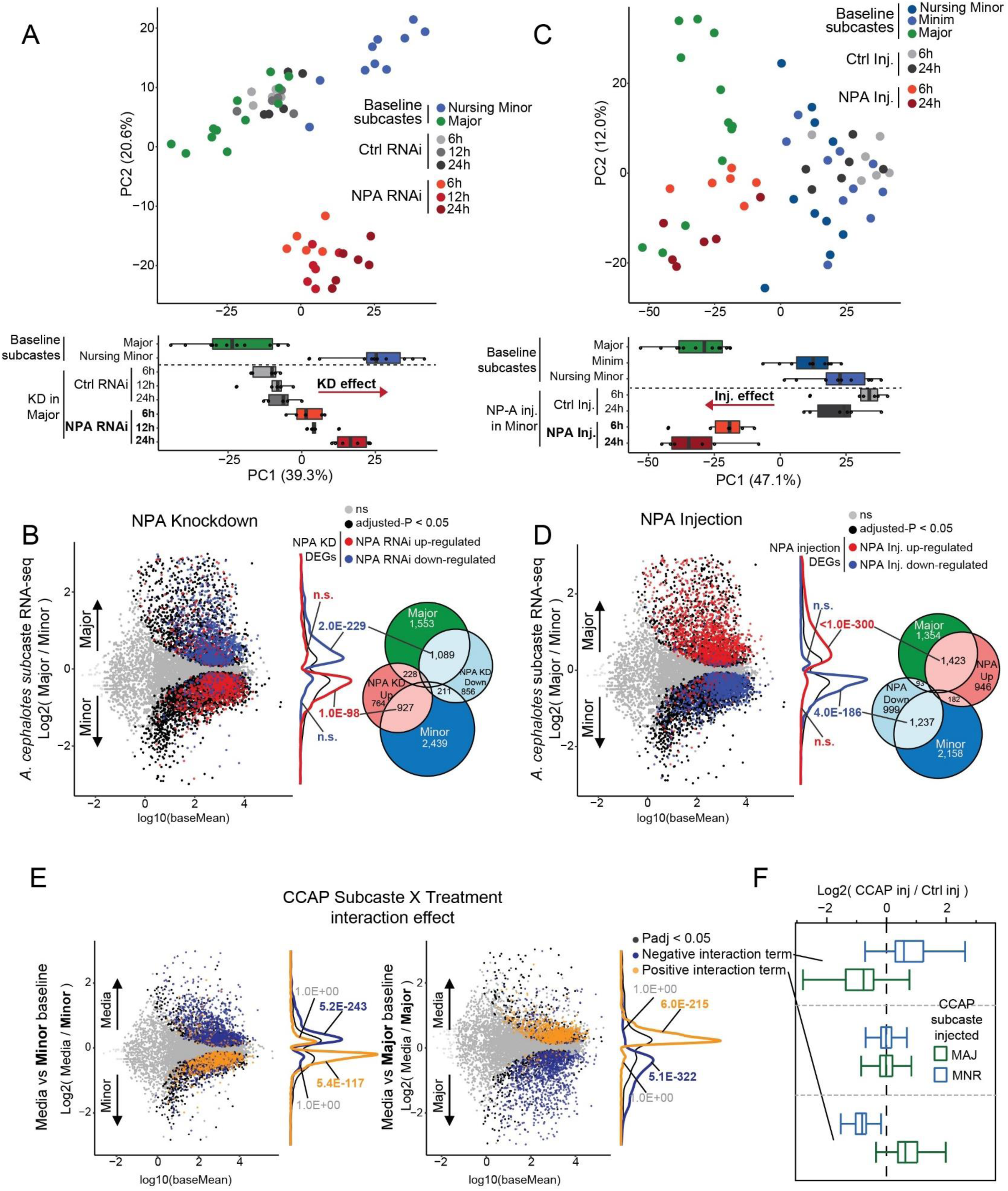
NPA manipulation illustrates centrality of NPA for Major worker expression. A) PCA of NPA KD. Subset to subcaste DEGs (as in Fig. 1B). Lower: values of each sample type along PC1, with arrow showing KD effect along PC1 comparing control to NPA KD. n=5-6. B) MA plot of Major/Minor baseline expression overlapped with NPA (or GFP control) KD DEGs (colored dots), illustrating stark transcriptomic reprogramming of Major workers following NPA KD. To the right is a venn diagram illustrating numbers of overlapping DEGs. P-values represent the results of a fisher’s exact test comparing groups. C) As for (panel A) but using NPA injection samples, illustrating striking effect of injection in shifting the Minor worker transcriptome from that resembling baseline Minor-like to baseline Major-like. Lower: values of each sample type along PC1, with an arrow showing injection effect along PC1 (as for panel A). n=6. D) The same as for (B) but utilizing NPA injection DEGs, illustrating reverse of panel B, where NPA injection reprograms Minor transcriptome. P-values represent the results of a fisher’s exact test comparing groups. E) CCAP interaction term from testing Major CCAP injection and Minor CCAP injection together (individual comparisons in Fig S5B), illustrating the strong opposite effect of CCAP injection depending on subcaste context. In orange are genes showing a positive interaction effect, which here, corresponds to genes that are upregulated in Major upon CCAP injection and downregulated in Minor upon CCAP injection. Blue represents genes showing a negative interaction effect corresponding to genes up-regulated in Minor upon CCAP injection and downregulated in Major upon CCAP injection. Left plots are (leftmost) differentially expressed genes between Media as compared to Minor and (middle) Media as compared to Major. Genes showing Minor up-regulation upon CCAP injection strongly overlap with genes biased to Media relative to Minor (left), and vice versa (middle). P-values represent the results of a fisher’s exact test comparing groups. n=6 for CCAP samples. F) Illustration of what significant interaction-effect genes shown in panel E represent, illustrating that negative interaction DEGs correspond to Minor up-regulated Major down-regulated (in CCAP injected samples relative to control injection) and positive corresponds to Major up-regulated Minor down-regulated (among CCAP-injected samples relative to control-injection).

Next, we compared DEGs following *NPA* knockdown in Major to genes significantly differing between baseline Major and Minor workers. Strikingly, we observed that genes up-regulated following *NPA* knockdown in Major strongly overlapped with genes biased to natural Minor, whereas down-regulated genes following *NPA* knockdown overlapped with genes biased to natural Major (**Fig. 4B**). We assessed *VKR* knockdown in Major and observed similar patterns among *VKR* knockdown DEGs shifting towards natural Minor along PC1, albeit less pronounced than for *NPA* knockdown (**Fig. S4B**, red arrow). Nevertheless, DEGs following *VKR* knockdown strongly overlapped subcaste DEGs similarly to *NPA* knockdown (**Fig. S4C**), consistent with the high correlation between overall transcriptional responses to *NPA* and *VKR* knockdown (**Fig. S4A**), and consistent with behavioral outcome (**Fig 3B**). These results show remarkable correlation between behavioral reprogramming of Major towards Minor caretaking, with transcriptional shifting of Major gene expression program towards natural Minor.

We performed RNA-seq of brains following injection of recombinant NPA peptide into Minors (see **Fig. 3D**), providing transcriptomics to match loss of caretaking behavior (see **Fig. 3D**) to contrast gain-of-function NPA knockdown in Majors (see **Fig. 3B, Fig S3I and G**). Indeed, DEGs of *NPA* knockdown in Major compared to NPA peptide injection in Minor were strikingly opposed in direction, with up-regulated genes in NPA injection in Minor corresponding to genes repressed by *NPA* knockdown in Major (and *vice versa*; **Fig. S4A**, right column). Further, PC analysis of NPA injection into Minor led to transcriptomic shift towards Major (**Fig. 4C**, blue shifting to red, towards green), which was opposite of *NPA* knockdown in Major shifting to Minor transcriptome (**Fig. 4B**). Genes up-regulated by NPA injection into Minor strongly overlapped with genes normally biased to baseline Major, and genes down-regulated by NPA injection strongly overlapped with genes normally biased to baseline Minor (**Fig. 4D**). These results show that loss of caretaking in Minor by NPA peptide injection leads to a transcriptomic shift toward Major, and this was opposite of gain-of-caretaking in Major by *NPA* knockdown.

We examined gene pathways in the behavioral reprogramming. Overall, we observed a strong degree of overlap between genes showing differential expression in Major, upon *NPA* knockdown or *VKR* knockdown, and following NPA injection in Minor: 253 genes in GO analysis showed lowest expression in Major, and up- regulation with both *NPA* and *VKR* knockdown in Major, and down-regulation upon NPA injection in Minor (**Fig. S4D,** right), and 403 genes showed the opposite pattern (**Fig. S4E,** right). Similar to functions delineating Major from other subcastes, genes showing highly consistent NPA repression were associated with neuronal, developmental, and ecdysone response terms (**Fig. S4D**, right), whereas genes showing NPA induction and Major-bias were strongly associated with metabolic and biosynthetic processes (**Fig. S4E,** right; **Table S5**).

Limiting these genes to those possessing a probable DNA-binding domain further supported the contrasting regulation: transcription factors of polycomb repression, brahma chromatin remodeling complex, and ecdysone response were identified among NPA-repressed genes (in natural Minor and reprogrammed Major) (**Fig. S4D,** left heatmap), whereas transcription factors with roles in regulation of metabolism and energy homeostasis were enriched among NPA-induced genes (in natural Major and reprogrammed Minor) (**Fig. S4E**, left heatmap). Taken together, the *NPA* knockdown and peptide injection experiments revealed that depletion of *NPA* shifts the Major transcriptome to that of Minor, underpinning the gain of nursing behavior (**Figs 3B, S3I and G**) and lowering of aggression (**Fig S3F**); in contrast, NPA injection shifts the Minor worker transcriptome to that of Major underpinning loss of nursing and increased aggression (**Fig S3F**). The *NPA*-dsiRNA-reduced genes were associated with a repression of chromatin remodelers and ecdysone response genes, while NPA- peptide-stimulated genes were associated with metabolic and biosynthetic functions and regulation.

We next evaluated whether CCAP perturbations show similar concurrence between behavioral (see **Fig. 3F-H, S3I and J**) and molecular outcomes. As mentioned above, CCAP peptide injection shifted behavior of both Major and Minor to natural leaf movement behavior of Media (see **Fig. 3F-G, S3I and J**). We injected CCAP peptide into brains of both Major and Minor followed by RNA-seq, hypothesizing that the response to CCAP peptide injection would differ between subcastes—which have very distinct transcriptomes at baseline (see **Fig. 1C-D**)—with the predicted outcome to shift both subcastes towards a common Media-like transcriptomic state. Indeed, PCA analysis of CCAP injected (compared to control peptide injected) brains showed that Minor shifted towards Media on PC1, while Major shifted in the opposite direction along PC1 (**Fig. S5A**, boxes and circles, respectively). Similarly, comparison of DEGs associated with CCAP injection showed opposite overlap with Media-biased expression, depending upon the subcaste injected (**Fig S5B, S5C**). Hence, both subcastes prominently converged towards the transcriptome of Media, but from opposite transcriptional directions depending upon the baseline transcriptome of the injected subcaste.

Because the outcome of CCAP injection varied transcriptomically by subcaste, we implemented a computational approach based on linear modelling to analyze both Major and Minor following injection. Here we utilized an interaction term to identify genes showing differences in their changes in gene expression following CCAP injection, depending on the subcaste of CCAP injection (see Methods^54^). This approach enabled identification of genes showing opposite changes upon CCAP injection, depending upon subcaste. We found sets of genes showing Major up-regulation and Minor down-regulation with CCAP injection (positive interaction effect) or Major down-regulation and Minor up-regulation with CCAP injection (negative interaction effect) (**Fig 4E-F**). This revealed numerous genes (>3000 per effect) showing significant interaction effect, indicating a striking difference in CCAP-dependent gene expression depending on subcaste-of-injection (**Fig. 4E-F**). One key observation was of genes showing Major up-regulation but Minor down-regulation upon CCAP injection, and these strongly overlapped with genes biased to Media (natural leaf movement) relative to Major (**Fig. 4E**, right, orange), while genes showing Minor up-regulation but Major down-regulation upon CCAP injection strongly overlapped with genes biased to Media (natural leaf movement) relative to Minor (**Fig. 4E**, left, blue). This was further underscored when examining genes showing bias to Media (as compared to Minor) and up-regulation upon CCAP injection into Minor (**Fig. S5D**), and *vice versa* (**Fig. S5E**). Notably, similar to NPA, CCAP injection into Minor resulted in up-regulation of genes associated with energy homeostasis and metabolism, including several transcription factors up-regulated upon NPA injection (e.g. *sug*, *bigmax*, *STUB1*; **Fig. S5D**, note GO analysis, lower), however CCAP injection into Major resulted in up-regulation of many genes associated with neuronal signaling, behavior, and vitellogenesis (**Fig. S6A**, note GO analysis, right), illustrating the striking transcriptomic reprogramming that occurs in Major upon CCAP injection.

Overall these findings show that injection of each neuropeptide into the opposite subcaste results in notable transcriptomic shift towards the subcaste which shows the highest baseline expression of the given neuropeptide. Thus, NPA injection shifted Minor to a Major-like transcriptomic state (**Figs. 4A-D**, **S4B**), whereas CCAP shifted both Major and Minor to a Media-like transcriptomic state (**Figs. 4E**, **S5A-B**). Importantly, the transcriptomic results were consistent in both neuropeptides with the behavioral outcomes following neuropeptide perturbation.

Because of the very strong impact of these neuropeptides on subcaste behavior and gene expression, we analyzed several neuropeptides or receptors we and others had previously observed to show clear subcaste bias. Examination of these revealed that many were regulated as predicted with NPA perturbation and CCAP injection relative to their baseline levels. Among several examples (**Fig. S5D**), *Ilp1*, a well-known insulin peptide which shows stark bias to Major^55^, showed strong up-regulation in Minor after NPA injection, showed repression in Majors after *NPA* knockdown, and opposite effects of CCAP injection depending on the subcaste injected (up-regulation in Minor and down-regulation in Major upon CCAP injection; **Fig. S6B**, left top).

Conversely, *sNPF-R* (the receptor for the neuropeptide sNPF, linked to regulating food intake and body size)^56^, which shows strong bias to Minor and Minim, showed the opposite pattern, with repression upon NPA injection, up-regulation 24h after *NPA* or *VKR* KD, and opposite patterns of differential expression with CCAP injection (**Fig. S6C**, top row). Collectively, we find that neuropeptide expression is strongly linked to focal behaviors in *A. cephalotes* (**Fig. 6**), and that these behavioral changes and reprogramming are paralleled at the transcriptomic level.

### Comparison with naked mole rat

To provide potential insight into behavior beyond the ant system, we compared subcaste-differential gene pathways in *A. cephalotes* to a eusocial mammalian species, *Heterocephalus glaber,* the Naked Mole Rat (*H. glaber*). *H. glaber* convergently possesses forager and non-foraging caretaking subcastes via age polyethism among workers, where young workers serve as nurse and older workers serve as forager^57^. In their natural state, foragers labor communally to dig a network of tunnels to find food, while nurses care for newborn pups by providing general protection and transferring fecal pellets and geophytes for nutrition.

We collected forager and nurse subcaste individuals from a mature *H. glaber* colony two weeks following the birth of a litter. This timeframe facilitated observation of foraging and nursing behaviors because the pups at this age require extensive care. To identify nurses, we measured interactions with pups, as defined by any instances of body-to-body contact, feeding the pups, and residence within the nest over a 20-minute interval (**Fig. S7A**, upper left and middle). To identify foragers, we quantified instances of food acquisition and time spent in tunnels distal from the nest (**Fig. S7A**, upper right). Given our interest in neuropeptides and functional orthologs within the pars intercerebralis in insects (where NPA and CCAP are localized and secreted), we performed subcaste-specific RNAseq on the *H. glaber* hypothalamus due to its role as a major neurosecretory organ/region that is functionally similar to the pars intercerebralis. We note that there have been few reported transcriptomic analyses of *H. glaber*^58–60^, and, to our knowledge, no reported transcriptomics of subcastes in eusocial mammals, and no reported comparisons between eusocial ants and eusocial mammals.

We compared genes differing in the hypothalamus between *H. glaber* forager and nurse to ant orthologs showing significant differences between in-nest and out-nest subcastes in *A. cephalotes* (e.g. see **Fig. 2A**; **Table S6**). This revealed remarkable overlap between *A. cephalotes* and *H. glaber* subcaste DEGs across >600 million years: genes more highly expressed in *A. cephalotes* out-nest subcastes (Major and Media) overlapped genes biased to *H. glaber* foragers, and similar overlap of in-nest *A. cephalotes* subcastes (Minor and Minim) with *H. glaber* nurses (**Fig. 5A** and **S7B**). Indeed, specific pairwise comparisons also showed overlap, with strong correspondence between genes biased to out-nest *A. cephalotes* subcastes Media (**Fig. 5B**, green genes) or Major (**Fig. S7C**, green genes) with genes biased to *H. glaber* foragers, whereas genes biased to Minor in-nest subcastes in *A. cephalotes* corresponded with genes biased to *H. glaber* nurses (**Fig. 5B** and **S7B-C**, blue genes). Further, comparisons involving Media vs Minor or Minim in *A. cephalotes* showed stronger correspondence with *H. glaber* subcastes (**Fig. 5B** and **S7D**) than those involving *A. cephalotes* Major vs Minor or Minim (**Fig. S7C**, **S7D**, bottom), however all comparisons showed similar trends and were significant. Comparison of specific genes in each subcaste (e.g. **Fig. S1B**, lower) to *H. glaber* subcaste-biased expression further supported this, with genes biased to Minor relative to all other subcastes showing the strongest association with *H. glaber* nurses and, similarly, Media association with *H. glaber* foragers (**Fig. S7E**). This underscores the correlation between foraging (the specific role of Media in *A. cephalotes* compared to Major defense) between species. Collectively, we show a remarkable overlap of subcaste-based orthologs of eusocial species despite divergence of >600 million years.

**Figure 5.**
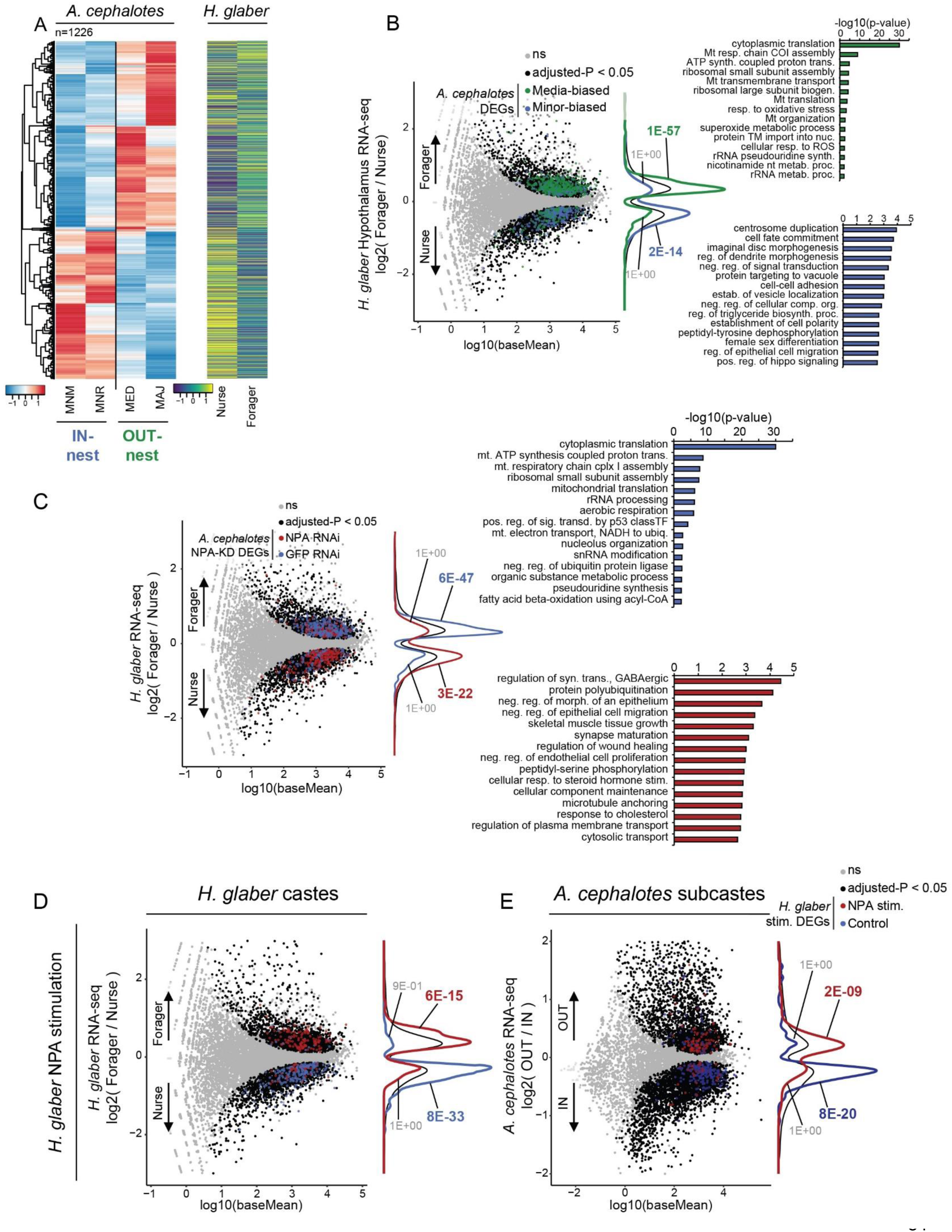
NMR shows correspondence in subcaste bias with *A. cephalotes* across 600my of evolution. A) Heatmap of *A. cephalotes* subcaste-biased gene expression for genes differing between the two major subcaste groups as compared to (right) gene expression among homologous genes from nurses and foragers of *H. glaber* for Hypothalamus samples (n=3 each sample type), showing overall correspondence in gene expression between species. B) MA plot of *H. glaber* gene expression differences between Nurse and Forager from Hypothalamus, overlapped with DEGs from *A. cephalotes* Minor (nursing morph) and Media (foraging morph) subcastes, showing significant overlap as shown in panel A. P-values represent the results of a fisher’s exact test comparing groups. Right: top GO terms (as determined in *H. glaber*) associated with each group of overlapping genes. C) Overlap between NPA KD DEGs (red and blue) and *H. glaber* hypothalamus differential expression. Right: gene ontology terms associated with genes showing the respective overlap (NPA KD down-regulation in *A. cephalotes* and forager-bias in *H. glaber*, and NPA KD up-regulation and nurse bias). P-values represent the results of a fisher’s exact test comparing groups. Right: top GO terms (as determined in *H. glaber*) associated with each group of overlapping genes. D) Overlap between *H. glaber* NPA stimulation DEGs (n=6, each sample type) with hypothalamus subcaste-biased expression as in B, showing NPA up-regulates genes enriched among forager-biased DEGs. P-values represent the results of a fisher’s exact test comparing groups. E) The same as for D but overlapping H. glaber NPA stimulation DEGs with A. cephalotes subcastes. For *A. cephalotes* MA plots all genes are shown but only genes with orthologs to H. glaber were used in significance testing. P-values represent the results of a fisher’s exact test comparing groups.

**Figure 6:** Schematic for leafcutter ant subcaste baseline behavior and reprogramming. Boxed are representations of each of the four subcastes’ baseline behavior focused on here. Upper: NPA-based interventions leading to suppression of caretaking behavior upon NPA injection into Minor workers and acquisition of caretaking behavior upon NPA KD in Major workers. Lower: CCAP interventions in Major and Minor workers both lead to the acquisition of Media-like leaf harvesting behavior.

Given these striking comparisons between subcastes, we returned to our targeted neuropeptides to investigate conservation in downstream foraging and nursing genes in *H. glaber* correlating with the two gain-of-function phenotypes with NPA and CCAP in *A. cephalotes*. We thus compared genes differentially expressed following *NPA* knockdown in *A. cephalotes* (acquisition of nursing in Major/Media, **Fig. 3B-C**) to genes differentially expressed between *H. glaber* subcastes. To examine nursing-related genes, we compared genes that were upregulated following *NPA* knockdown in *A. cephalotes* (associated with Major acquisition of brood care behavior) with genes biased to *H. glaber* nurses (**Fig. 5C**, red genes); these overlapping genes were enriched for multiple terms related to neurodevelopment and general developmental processes (**Fig. 5C**, right red GO terms). In contrast, to examine foraging-related genes, we compared genes that were down-regulated following *NPA* knockdown in *A. cephalotes* with genes more highly expressed in foragers in *H. glaber* (**Fig. 5C**, blue genes and right blue GO terms). Specifically, we observed strong correspondence between genes that decrease in expression following *NPA* knockdown with those more highly expressed in foragers in *H. glaber*, and vice versa (**Fig. 5C**). Genes showing decreased expression upon NPA knockdown (i.e. NPA up-regulated genes) in *A. cephalotes* that were also biased to *H. glaber* foragers showed similar enrichment of functional terms as those generally biased to foragers; these include enrichment of translation and mitochondrial terms, and other metabolism-related functions (**Fig. 5C**, right). Genes showing up-regulation after NPA knockdown in *A. cephalotes* (i.e. NPA down-regulated genes) that were also biased to *H. glaber* nurses were enriched for multiple terms related to neurodevelopment and general developmental processes (**Fig. 5C**, right). We performed the same analysis using genes responsive to CCAP injection in Minor and found a similar pattern, with genes up-regulated following CCAP injection overlapping with genes biased to *H. glaber* foragers, and vice versa (**Fig. S7E**). Functions associated with each class of overlapping genes were similar with NPA responsive genes (**Fig. S7F,** right). These results broadly link gene expression between *A. cephalotes* and *H. glaber* subcastes and reveal that NPA and CCAP regulate many genes showing convergent evolution of subcaste-bias between ant and mammal eusociality.

We then investigated whether ant NPA can function in a mammalian species lacking it and lacking its receptor. We utilized cells in culture (see Methods) derived either from *A. cephalotes*^40^ or from *H. glaber* brain tissue and treated either culture with recombinant *A. cephalotes* NPA. First, we validated the use of the NPA in brain cells *in vitro* using the homologous ant system (NPA in the *A. cephalotes* neuronal culture). As anticipated, we found NPA up-regulated genes significantly overlapping with genes biased to *A. cephalotes* out-nest Major, and NPA-repressed genes more strongly overlapping with genes biased to in-nest Minor (**Fig. S7G**). Importantly, these results underscore the centrality of NPA in suppressing caretaking; specifically, the strongest pattern was NPA repressing genes normally biased to nurses (**Fig. S7H**), consistent with the behavior results of *NPA* knockdown leading to the acquisition of caretaking behavior (**Fig. 3A-E**).

Having established the efficacy of NPA treatment *in vitro* using ant cells, we then treated *H. glaber* astrocyte cultures (derived from the brain cortex of newborn pups) with NPA, and we compared altered genes *in vitro* to those differentially expressed between *H. glaber* nurse and forager. Strikingly, we found that genes up-regulated by NPA in *H. glaber* culture were enriched among *H. glaber* foragers (**Fig. 5D**, top; **Fig. S7H**, top), whereas genes down-regulated by NPA in *H. glaber* culture were associated with *H. glaber* nurses (**Fig. 5D**, bottom; **Fig. S7H**, bottom). These findings indicate that NPA was able to repress genes associated with nursing in *H. glaber* via a potentially conserved mechanism. Hence, we then compared genes across species, examining differentially regulated genes upon stimulation of *H. glaber* astrocytes with NPA, to genes differentially expressed between *A. cephalotes* subcastes. Strikingly, despite >600 million years of divergence, we found that these same patterns were maintained, with NPA-upregulated genes in *H. glaber* cells showing enrichment among *A. cephalotes* Major/Media workers (**Fig. 5E**), that is, despite being absent from *H. glaber*, NPA regulates an overlapping set of conserved genes associated with caretaking in both species.

One potential mechanism whereby NPA may mediate behavioral changes in a species entirely lacking it and its receptor is via partial agonism of insulin signaling, as suggested in a prior study^61^. Because of this, we returned to our caretaking behavioral assay in *A. cephalotes* (see **Fig. 3A**) but this time injected insulin or the typical vehicle control^62^ (BSA) into Minor workers. This resulted in a similar behavioral effect as for NPA, with reduced brood movement in Minors injected with insulin (**Fig. S7I**, n = 32, p-value = 1.1E-02), indicating insulin peptide injection is able to phenocopy NPA peptide injection in *A. cephalotes*. This is consistent with prior reports linking NPA to insulin signaling^32^ and may suggest NPA is able to tap into insulin signaling even across 600 my. Overall, these results illustrate surprising conservation in molecular underpinnings of caretaking behavior, despite the emergence of such behavior occurring independently in these species and NPA being absent from mammals.

## DISCUSSION

Here we report extensive molecular characterization of leafcutter ant worker subcastes and of eusocial naked mole-rat non-breeder subcastes (equivalent to social insect workers), and the first comparison between them relevant to behavioral regulation by neuropeptides. To establish a rigorous system of testing the complex behavior of leafcutter ants, we utilized the species *Atta cephalotes* (*A. cephalotes*) and segmented worker subcastes into four morphological groups (Major, Media, Minor, and Minim) exhibiting distinct behaviors (**Fig. 1A**). Transcriptomic analyses of *A. cephalotes* worker subcastes revealed high-level similarities with signatures of caste in species with less morphologically-elaborated caste systems, likely related to differences in hormonal signaling (**Fig 1F, S1E**), however this was insufficient to explain differences between individual subcastes. Given that division of labor in *A. cephalotes* is based on worker neuroanatomical phenotypes that correlate with cognitive demands for task performance^36^, we investigated connections between levels of certain neuropeptides and the more elaborate number of *A. cephalotes* worker subcastes, via neuropeptide perturbation and examining subcaste-biased behavior and altered gene expression.

We focused on NPA and CCAP neuropeptides, which show, respectively, highest expression in Major and Media subcastes, as targets for perturbation. We demonstrated that NPA, which showed highest expression in Major workers (**Fig 2B**), as a potent regulator of brood care—a critical behavior distinguishing worker sub-castes and, more broadly, eusocial division of labor. Knockdown of *NPA* (or its putative receptor VKR) led to acquisition of brood care behavior in two subcastes (Major and Media) which never naturally engage in caretaking behavior (**Fig 3, S3**), while injection of recombinant NPA resulted in a blocking of brood care behavior in Minor (**Fig 3D**) and also increasing aggression (**Fig S3F**). We investigated a second neuropeptide, CCAP, in governing leaf “harvesting and transport”—a signature behavior of the leafcutter ants within the tribe *Pheidolini*^63^—due to its considerable elevation in Media workers (**Fig 2B**). Indeed, injection of CCAP peptide into Majors and Minors (castes with low natural expression of *CCAP* at baseline) resulted in acquisition of leaf moving behavior (**Fig 3F-G, S3I, J**), while *CCAP* knockdown in Media (caste with high natural CCAP levels) resulted in loss of baseline leaf movement (**Fig 3H**). Hence these neuropeptides have a profound effect on behavior, and are able to dominantly reprogram behavior of Major workers to caretaking or to leaf transport— behaviors that Majors never typically perform.

Transcriptomic profiling following perturbation of these neuropeptides showed consistency with the behavioral results: (1) *NPA* (or VKR) knockdown in Major led to acquisition of a transcriptomic state resembling the natural NPA-low in-nest Minor subcaste (**Fig. 4A, S4B**), (2) injection of NPA into Minor (natural brood care) led to a transcriptomic state resembling NPA-high Major (**Fig. 4C**), and (3) CCAP injection resulted in a shift of Major and Minor CCAP-low subcastes to a Media-like CCAP-high state (**Fig. 4E-F**). The results from CCAP were particularly striking, as injection of CCAP into Major or Minor resulted in many caste-specific genes biased to one or the other caste changing in direction toward the Media transcriptome, illustrating strong context dependence of CCAP action to accomplish the same behavioral outcome in distinct caste states. The mechanisms underlying CCAP ability to sculpt the transcriptome from either Major or Minor toward Media is a promising future avenue of investigation.

Together, these results illustrate the importance of neuropeptides to behavioral programming, particularly in the context of the elaborated worker subcaste system in *A. cephalotes.* We found that knockdown of NPA in *C. floridanus* Majors resulted in similar behavioral outcomes (**Fig 3E**), suggesting that, despite the differences in behavior between Majors of *A. cephalotes* and *C. floridanus* (*C. floridanus* Majors rarely leave the nest), brood care behavior is blocked by NPA in Majors of both species. Thus, the basis of NPA’s regulation of brood care behavior may be more basal than the evolution of Attine worker systems.

In support of this, across >600 million years of evolutionary divergence, DEGs resulting from neuropeptide perturbation in ant brain—targeting a neuropeptide only present in insects—showed strong, consistent overlap with *H. glaber* DEGs in both cases (**Fig. 5C, S7F-H**). More generally, we found an unexpected overlap in DEGs biased between *H. glaber* non-breeder (worker) subcastes and *A. cephalotes* baseline subcastes, suggesting convergence on similar pathways in the regulation of non-reproductive behavior (**Fig 5A-B**).

More specifically, our results indicate that the insect neuropeptide NPA alters expression of conserved genes that underlie caretaking behavior in both ant and mammal. Prior observations suggest that NPA may tap into the highly-conserved insulin signaling pathway^61^ which is supported by our results (**Fig S7I**), and consistent with functional terms related to conserved caste DEGs (**Fig 5B**) or NPA sensitive genes (**Fig 5C**). To our knowledge, insulin has not been previously linked to brood care in either species but has been associated with caste in social insects^64,65^. Further, in mammals insulin/IGF1 signaling has been linked to differences in anxiety-like behaviors^66^, memory formation^67^, and the similar ligand IGF1 has been posited to play a role in modulating maternal behaviors^68^. Thus, it is possible that insulin signaling has been converged upon in *H. glaber* for the regulation of non-reproductive caste behavior and would explain how NPA can lead to worker subcaste-associated transcriptomic changes in *H. glaber in vitro* despite natural NPA expression being confined to arthropods^69^. This may also suggest a role for insulin/IGF1 signaling more broadly in mammalian maternal caretaking behavior, as a key prediction in the evolution of eusociality is that sterile worker behavior is derived from ancestral maternal care behaviors^70^.

The identification of two neuropeptides challenges the prevailing notion that ant worker behavior is solely the result of developmental cues and emergent interactions between individuals in the colony. Traditionally, eusociality is attributed to complex social behavior and collective decision-making, controlled by external factors and communication between workers. However, the ability to modify behavior via neuropeptide pathways in adult ants suggest that physiological pathways are crucial in maintenance of worker behavior.

These findings suggest that eusociality may not only emerge from social dynamics. If worker behavior is easily reprogrammable, it suggests a higher potential for flexibility and adaptability than previously thought. This flexibility could be attributed to eusocial species needing to rapidly adjust to environmental changes without requiring genetic evolution across generations.

Collectively, our results further establish ants as a remarkable system to elucidate the foundations and maintenance of social hierarchies and offers molecular characterization of a species exhibiting extreme division of labor. Future investigation will reveal how NPA functions during leafcutter ant development including the establishment of caste and subcaste identity and plasticity of social states. Further, the many other differentially expressed genes and neuropeptides in our study offer abundant opportunity to deeply probe the evolution of division of labor across an expansive range of behaviors.

### Limitations of the Study

One limitation of this study is that the age of ants used here were not rigorously controlled. In this species we utilized behavior-matched individuals due to the difficulty of painting newly eclosed ants, and lack of persistence of any paint mark added to newly eclosed workers. Workers of most ant species exhibit age polyethism (changes in behavior associated with changes in worker age), and this is a complexity that we did not evaluate. In our previous studies in *C. floridanus*, we found that there is a vulnerable time window, after eclosion, to reprogramming that closes at 10 days^7^. While we found *A. cephalotes* workers susceptible to reprogramming even when using samples without age matching, it is possible reprogramming would be stronger at a specific age. Alternatively, the impact of age on behavioral reprogramming may be more limited than for *C. floridanus* due to differences in life history between the two species, however we did not assess this in this study.

A second limitation is the lack of clear homology of our focal neuropeptides in *H. glaber*. While our results suggest that similar transcriptomic changes in *H. glaber* brain cultures as *A. cephalotes* following addition of NPA may be due to insulin signaling, we did not test this pathway directly, and thus represents an open question in our study.

A final limitation is that of the resolution of our perturbations and analyses. While our identification and perturbation of neuropeptides at the whole brain level was impactful, this does not reflect the complexity of neurohormonal signaling and neural circuitry. It is likely that the behavioral outcomes here are due to more subtle changes than whole-brain rewiring, and thus, future studies are needed to focus on finer-scale analyses of baseline differences in neural circuitry between worker subcastes, and how these neuropeptides impact circuits regulating the behaviors assayed here.

## METHODS

### Atta cephalotes ants

Newly mated *Atta cephalotes* queens were caught in 2018 in Trinidad and raised for 2 years by Andrew Stephenson (vendor) in Scotland for 2 years before 3 genetically distinct colonies containing a single queen and about 5000 workers were brought to Berger lab. Each colony was housed separately, and colony effects were considered during experimental design (equal control and treatment from each colony) and statistics (blocking for colony effect). Each *Atta* colony was housed in a 50-gallon tank with fungus gardens and foraging chamber suspended above heated water to maintain a controlled temperature of 30°C and 70% humidity within our ant facility on a 12-hour light/dark cycle. The foraging chamber and fungus gardens were placed at equal heights to facilitate easy movement within the colony. The water was changed every 2 weeks to maintain over health of the colony. A Dremel was used to create slits in the bottom of the foraging chamber and fungus boxes to avoid buildup of condensation. Dirt was given to the ants weekly to plug air holes and maintain the humidity of the fungus gardens at 95%. Fungus health was monitored daily based upon on color, texture, and smell. The unhealthy fungus was removed if necessary. The ants were intermittently checked for mites and infections. The colonies were provided with leaves between 9-10am each day and example species include wild rose, privet, honeysuckle, oak, porcelain berry, chokeberry, and linden. The same species of were frozen and stored at −20°C to provide to the colonies during times when fresh vegetation was not available.

### Heterocephalus glaber housing and subcaste determination

All naked mole-rat experiments were approved and performed in accordance with guidelines instructed by the University of Rochester Committee on Animal Resources with protocol 2009-054. Naked mole-rats were originally caught in Kenya and transferred to the University of Rochester. For this study, we used two independent colonies called colonies 7-1 and 7-5, with housing conditions as described here: Briefly the animals are maintained in custom made chambers connected by plexiglass tubes to recreate the structure of underground borrows. A large central chamber is treated by the animals as a nest, evidenced by breeder and pup residence and colony member sleeping location. Distal chambers (connected by plexiglass tubes) where food is deposited (3 times per week) is treated as outside the nest and is only frequented by older non-breeders (foragers). Nurses were chosen based on instances of body-to-body contact and feeding the pups while foragers were chosen based upon food acquisition behavior and residence in distal tunnels. Sources of animals used in this study were as previously described^71,72^.

### Heterocephalus glaber tissue harvest

Naked mole-rat workers were behavior-matched and designated either forager or nurse. Designated animals were removed from the colony to small hand warmers beneath the cage to ensure the animal stayed warm. Body weight was measured, and animals were brought into fume hood in an induction chamber with isoflurane and 5% oxygen at 1L/min. We waited until the animal had shallow breathing and no toe reflex. The animal was removed from the induction chamber and placed on its back on a clean Styrofoam board. A nose cone was placed over the animal’s head with isoflurane and equivalent oxygen concentration. Two front and two back feet were pinned on Styrofoam. The chest cavity was opened and the rib cage was held back with a hemostat. A 25g needle and 1cc syringe was inserted bevel-side up into the right ventricle of the heart to remove as much blood as possible (0.2-0.6 mL) to avoid cross-contamination of innervated tissues. The brain was dissected and the hypothalamus and hippocampus were removed for RNA-seq. At least one sample was taken from each of two colony backgrounds (7-1 and 7-5) to avoid pseudoreplication.

### Cell culture

Neonatal naked mole-rat cells were extracted from newborn pups (n=5). Astrocytes were extracted from brain cortex using neural tissue dissociation kit (Miltenyi Biotech, cat. 130-092-628) according to the manufacturer’s instructions. The cells were cultured in DMEM/F-12 medium containing 10% fetal bovine serum (GIBCO). When cells were 80% confluent, isolated cells were frozen in liquid nitrogen within 2 passages. All subsequent cultures were performed in EMEM (ATCC, 30-2003) supplemented with 15% fetal bovine serum (GIBCO), 100 units/mL penicillin, and 100 mg/mL streptomycin (GIBCO). All primary cells were cultured at 37C with 5% CO2 and 3% O2.

Naked mole-rat astrocytes (n=1X10^5^) were seeded in 6-well plates. The next day cells media was replaced with the new media containing one of the following reagents: 10 μg/ml neuroparsin (cat. Vendor) or 10 μg/ml scramble peptide (cat. Vendor). This neuropeptide does not share significant homology with any naked mole-rat genes but maintains putative sites to bind to receptors involved in insulin signaling^73^. The cells were washed in PBS and harvested 6h later in Trizol reagent (Thermo Scientific, cat. 15596026).

### Peptide expression in Sf9 cells

Sf9 cells (Invitrogen, #11496015) were expanded in Sf900 III medium to passage 3 and cryopreserved at −120°C. Cryopreserved passage 3 cells were thawed and expanded to passage 6 and then cryopreserved at −120°C. Sf9 cells were maintained as suspension cultures in shake flasks and grown in Sf900-III medium (Invitrogen, #12658035) at 27°C at 160 RPM. Cells were seeded 3 times per week (usually Monday, Wednesday, Friday) at a density of 0.5-1.0 x 10^6^ cells/ml to maintain cultures in logarithmic growth. Cells were maintained for a maximum of 30 passages. Monolayers of Sf9 cells cultured in antibiotic free SF900III medium were transfected with Cellfectin (Invitrogen) and a PCR validated, recombinant bacmid DNA constructed by transposition of the donor plasmid DNA in *E. coli* cells, the so-called Bac-to-Bac™ (Invitrogen-Gibco/Life Technologies) method. The source of genomic baculovirus DNA was from *Autographica californica nuclear polyhedrosis* virus (AcNPV). The tissue culture supernatant from transfected cells was harvested 120 hours post-transfection and considered the P1 virus stock, which typically has a titer in the 5 x 10^6^ pfu/ml. High titer P2 virus stock was produced from the P1 virus (the original virus from cell culture supernatant of transfected cells) in Sf9 cells grown in SF900 III medium as a suspension culture in shake flasks. A high-titer P2/P3 virus stock was produced by infecting Sf9 cells with a virus stock at a low multiplicity of infection (MOI=0.1) and culturing for 120 hours. Typical titers were 1-4 x 10^8^ pfu/ml. A 100 ml suspension culture of Sf9 cells grown in SF900-III were infected with a high-titer baculovirus stock at an MOI=1-2. Cells or conditioned media (for secreted proteins) were harvested at 24-, 48-, and 72-hours post infection to evaluate the integrity, stability, solubility, and optimum yield of the desired product(s). High density (2 x10^6^ cells/ml) suspension cultures of Sf9 cells grown in Sf900 III were infected at a multiplicity of infection (MOI) equal to 1-2 for 48-72 hours. Infected cells were harvested by centrifugation at 500xg for 5 min.

### Peptide purification

Ammonium sulfate was added to the Sf9 media to 60% saturation. The precipitant was then filtered to separate the media components. Then the precipitant was resuspended in 80 mL of lysis buffer (50 mM HEPES pH = 7.4, 150 mM NaCl, 2 mM Beta-mercaptoethanol, 5% glycerol) per 1L of precipitated Sf9 media. 50 mL of lysis buffer was then used to equilibrate 5 mL of FLAG resin and the resin was incubated in the resuspension for 1 hour at 4°C with gentle shaking. The mixture was then poured over a gravity column to allow the supernatant to run over the resin. The resin was then washed twice with 25 mL of lysis buffer. The resin was then incubated with 10 mL of lysis buffer supplemented with 300 μg/mL of 3X FLAG peptide at room temperature with gentle shaking and eluted by running over the gravity column. The elution was concentrated in a 3k MWCO spin concentrator to 500 μL. Then we equilibrated an S75 column with SEC buffer (50 mM HEPES pH = 7.2, 150 mM NaCl, 2 mM TCEP, 5% glycerol). Then we ran the concentrated elution on the S75 column and peaks were analyzed on an SDS-PAGE gel.

### Peptide injections

Minors were removed from the main colony and ants received brain injections with either purified neuroparsin-A or a scramble peptide control. Failed injections (death within 24 hours) were excluded from the study, but rigorously recorded. The mortality rate of injected Minors never reached above 5% in both conditions up to 1 week post-injection. The success of the injection was also verified by daily checks of the ants’ mobility, which remained similar to uninjected ants of the same subcaste. The scramble peptide allowed investigation of behavior changes as a function of the injection. Injections were performed with a robotic arm holding a borosilicate needle filled with the peptide to the injection site (front of head between the antennae), and a Femtojet micro-injector to inject 100 ng peptide dissolved in 0.5 μL of 1x HBSS. CCAP (Thermo Scientific - HY-P0303) or scramble peptide (custom synthesis from Thermo) were injected into either Majors or Minors. Insulin peptide (Thermo – 12585014) was injected into Minors using the same method described above at 100 ng per injection. This insulin has 53.8% homology with insulin-like peptide 1 (Ilp1) in *Atta cephalotes* – the most differential insulin peptide between worker subcastes. NPA has a 34.1% homology with IGFBP7 in *Homo sapiens*, which can bind the insulin receptor. NPA does not share significant homology with *Homo sapiens* insulin, but shares structural residues and may function similarly^73^.

### Mass Spectrometry

Ant brains were lysed in 50 μL of a buffer of 8 M urea, 100 mM NaCl, 50 mL Tris-HCl (pH 8) and an inhibitor cocktail (Thermo Fisher Scientific, Halt Protease and Phosphatase Inhibitor Cocktail #78442). Brains were subjected to 2 x 30 s of sonication with a Bioruptor (Diagenode), 20 min of incubation on ice, followed by 10 min of centrifugation at 21,000 × g and 4°C. Protein content of the supernatant was estimated with using BCA. The proteins were reduced with 5 mM dithiothreitol for 30 min at 55°C and alkylated with 10 mM iodoacetamide in the dark for 30 min at room temperature. The proteins were diluted to 2 M urea with Tris-HCl and digested with sequencing-grade trypsin (Promega) at an enzyme:protein ratio of 1:25 for 12 hr at 37°C. The digestion was quenched with 1% trifluoroacetic acid. Samples were then desalted using Pierce C18 Tips, 10 µL bed (Thermo Scientific). Peptides were brought to 1 μg/3 μL in 0.1% formic acid (buffer A) prior to mass spectrometry acquisition. To build the chromatogram library, aliquots from each biological replicate from each subcaste were pooled to ensure that the pool contained virtually every peptide present.

### Immunoprecipitation-Mass Spectrometry of VKR

5 μg of FLAG-NPA was injected into Major heads and the ants were placed in a recovery chamber for 1 hour. The recovery chamber was a clear plastic container (Pioneer Plastics – 028C) with the bottom covered in hydrated gypsum cement to maintain high humidity. The walls were coated with Polytetrafluoroethylene (Thermo Cat# 665800) to prevent the ants from escaping. The recovery chamber also included fungus as a food source. The brains were dissected following recovery and protein lysates were prepped. 135 μL of protein A and 135 μL of protein G were washed with 1 mL of 0.5% blocking buffer and rotated for 10 minutes at 4°C. The beads were bound to a magnetic stand and 3 washes were performed with 1 mL of the 0.5% blocking buffer. After the final wash, the beads were bound to the magnetic stand and the washing solution was removed. Beads were resuspended in FLAG antibody in the blocking buffer. Antibody was incubated for 5 hours and then protein lysate pooled from 20 Major brains was added for each IP. Sampled were for mass spectrometry with Protifi S-trap micro columns and desalted with C18 columns. The samples were run according to the bottom-up mass spectrometry method described below.

### Bottom-up nanoLC-MS/MS and data analysis

Samples were analyzed in 6 biological replicates per subcaste with nanoLC-MS/MS. Peptides were separated using an UltiMate3000 (Dionex) HPLC system (Thermo Fisher Scientific, San Jose, CA, USA) using a 75 μm ID fused capillary pulled in-house and packed with 2.4 μm ReproSil-Pur C18 beads to 20 cm. The HPLC gradient was 0%–35% solvent B (A = 0.1% formic acid, B = 95% acetonitrile, 0.1% formic acid) over 90 min and from 45% to 95% solvent B in 30 min at a flow rate of 300nl/min. The QExactive HF-X (Thermo Fisher Scientific, San Jose, CA, USA) mass spectrometer was configured following a data-independent acquisition (DIA) chromatogram library method^74^. For wide-window data, full scan MS spectra (m/z 385-1015) were acquired with a resolution of 60,000, AGC target of 1e6; MS/MS spectra were acquired with 8 m/z staggered isolation windows (normalized collision energy of 27.5 and an AGC target of 1e6). The chromatogram library was collected in 6 gas-phase fractions (GPF) with DIA with full scan MS spectra of 110 m/z each with 4 m/z overlapping windows with the same resolution and AGC targets settings as the wide window data. Mass spectrometry data files were demultiplexed with MSConvert^75^. Chromatogram library was built from 6 gas-phase fractions using Walnut in EncyclopeDIA. Wide-window data was processed in EncyclopeDIA with a chromatogram library from Walnut. Processed mass spectrometry data files were imported into Skyline. For statistical analysis, Skyline data was exported to MSstats, normalized by equalizing medians, and a 2-tailed t-test was performed (significant if adjusted-p < 0.05)^76^.

### Behavioral assays

For behavioral assessment we performed both automated and manual assessments of perturbation samples. This was due to simplicity of behavioral outcomes following perturbation, however automated tracking revealed similar results. For both peptide and dsiRNA injection the head capsule of ants was injected, followed by recovery for 1h in a small chamber (Pioneer Plastics – 028C), followed by transfer to the final behavioral chamber for assessment, which was begun 1h after transfer to allow for acclimatization to the behavioral arena.

#### For automated assessment of perturbations

(**Fig S3G, I, J**), as well as behavioral analysis of baseline castes (**Fig 1B**) videos of ants (Fig. 1B) were analyzed using DeepLabCut^77^ (version 2.2b8), a marker-less pose-detecting machine learning Python package. This toolbox utilizes Tensorflow and ResNet to track any user-defined body part, animal, or object in successive video frames. DeepLabCut is blinded to experimental conditions and is capable of tracking movements that allow quantification and analysis of locomotion and therefore behavioral differences. We utilized an Exxact workstation with the specifications: Valence VWS-1542881-DPN-Deep Learning DevBox (X299, 1x i9-9820X, 16GBx8 DDR4, 1x 2TB NVMe OS SSD, Dual 10GBase-T Onboard, 2x Titan RTX 24GB GPU with NVLINBK Bridge, Ubuntu 18.04). We trained the neural network with 1,000,000 iterations on 60 labeled frames. We labeled 51 individual body parts on each ant. This regimen was adequate to produce the desired fit of the model to the training data (loss <0.005), such that prediction of movements by DeepLabCut resulted in accurate tracking of body parts. For actual assessments we utilized the thorax mid-point and head mid-point body parts exclusively to score distance to/interaction with assay objects, as well as locomotion. For interaction with assay objects, an ant was required to fall within 25 frames of a given object (fungus, brood, leaves) and both the ant component and assay object were required to have a likelihood of >0.8. Proportions of frames spent near a given object were then assessed. For Locomotion, average distance per-frame was calculated across the measurement period.

#### For manual scoring of perturbation behavior

All assessments were investigator blinded. For the brood-movement assay, 1 gram of fungus from the parent colony was placed in the upper left impression of the behavior chamber. Four pupae were placed in the bottom impression and 10 grams of leaves were placed in the right impression of the behavior chamber. The chamber was washed with pentane between experiments to remove residual cuticular hydrocarbons (pheromones). We defined a “brood movement” as picking up a pupa from the bottom impression and moving it to the fungus impression. 20-minute videos were taken of ants in the behavior chamber using a Nikon camera equipped with an AF-S DX Micro NIKKOR 40mm f/2.8G (30 frames per second). For NPA perturbations, brood movement was counted when the ant fully transported a pupa from its original chamber to the fungus area. For CCAP perturbations, leaf movement was counted when an ant moved a piece of leaf (pre-cut) outside the leaf chamber to the rest of the behavior arena. For CCAP perturbations, pre-cut leaves were used as we were unable to coax the workers (including the untreated leaf cutting subcaste, Media) to cut leaves in the context of our artificial assay, so we thus used leaf harvesting/collecting as a proxy for Media worker behavior.

### Brain cell count

For each sample 3 freshly dissected brains were transferred to 200 μL of cold nuclei isolation buffer (10 mM Tris-HCl, pH = 7.4), 10 mM NaCl, 3 mM MgCl2, 1% BSA, 0.1% Tween-20, 0.1% NP-40), and dissociated with a 0.5 mL Dounce homogenizer (25 strokes with pestle A and 40 strokes with pestle B). The homogenized sample was transferred to a 1.5 mL Eppendorf tube with 1 mL of pre-chilled washing buffer (10 mM Tris-HCl, pH = 7.4, 10 mM NaCl, 3 mM MgCl2, 1% BSA, 0.1% Tween-20) and incubated on ice for 5 minutes. Nuclei were pelleted using a swinging bucket centrifuge for 10 minutes at 500 g to minimize sample loss. The pellet was resuspended in 200 μL of DPBS and DAPI for nuclei counting in a hemocytometer under a fluorescence microscope. The brain cell number for each subcaste was calculated based on the nuclei counting normalized to input volume and brain number.

### RNAi Injections for Neuroparsin-A and VKR

Custom short interfering RNAs (DsiRNAs) targeting Neuroparsin-A and VKR were synthesized by IDT. DsiRNAs were resuspended to 20μM. 10 μL aliquots were taken and mixed with 10 μL of 10% glucose, 1.5 μL of in vivo-jetPEI transfection reagent (Polyplus Transfection) and 8.5 μL of Milli-Q water. The solution was incubated at room temperature for 15 minutes prior to injection. Each ant was injected with a calibrated glass capillary needle directly into the head. 2 μL was injected for Majors and 0.5 μL was injected for Minors.

### RNA-seq sample selection, RNA extraction, and library preparation

For baseline caste RNA-seq sample selection, workers were first assessed based upon morphology (**Fig S1A**), and then among a given morphometric group (**Fig 1A**), only samples exhibiting a specific behavior were selected for downstream RNA-seq: Minim, interacting with fungus and hitchhiking; Minor, brood interaction, leaf processing, and midden work; Media, leaf cutting and carrying of leaves; and Major patrolling outside the nest. Following analyses using each subcaste X behavior (eg Fig S1C, top) and finding more blunted transcriptomic differentiation when using these segregations, we performed comparisons involving baseline castes (eg Figs 1C-F, S1C-E, 2A-C, S2A-B) using ‘gardening’ Minims, nursing/brood-associated Minors, leaf carrying+cutting Media, and all Majors for establishing DEGs, while still plotting all samples.

*A. cephalotes* whole brains were dissected from single individuals, selected as above, snap-frozen and then homogenized in TriPure (Sigma), followed by precipitation in isopropanol. RNA was then subjected to DNase treatment in 25uL, followed by re-purification via RNAclean XP beads (Beckman Coulter) using a 2:1 ratio of beads:sample. For RNA-seq, polyadenylated RNA was purified from total RNA using the NEBNext® Poly(A) mRNA Magnetic Isolation Module (NEB E7490) with on-bead fragmentation as described^88^. cDNA libraries were prepared the same day using the NEBNext® Ultra™ II Directional RNA Library Prep Kit for Illumina® (NEB E7760). Following adapter ligation and purification qPCR was used to determine optimal cycle numbers for library PCR, and all samples for a batch were run the number of cycles determined for the lowest-quantity sample in order to avoid sample-specific biases introduced by PCR cycle number differences.

### Assignment of gene orthology and functional terms

Genes (*Atta cephalotes* NCBI RefSeq annotation release 100; assembly vGCF_000143395) were assigned orthology using the reciprocal best BLAST hit method^89^ to both *D. melanogaster* (r6.42) and *H. sapiens* (GRCh38) protein coding genes.

GO terms were assigned to genes in the following way: GO terms from reciprocal best hit *D. melanogaster* orthologs were lifted over to the corresponding genes. These were supplemented with GO terms assigned by running interproscan on the entire geneset and collapsed to a non-redundant set. For genes lacking a reciprocal best-hit ortholog in *D. melanogaster*, standard one-way blast hits were used to assign terms from *D. melanogaster* (e-value cutoff: 1E-15). For *H. glaber* the same approach was taken as above but using *M. musculus* instead of *D. melanogaster*.

GO enrichment tests were performed with the R package topGO^78^, utilizing Fisher’s elimination method, and resulting significant terms were entered into REVIGO^79^ for collapsing of redundant terms.

### A. cephalotes annotation update

Because we observed a large number of genes which possessed truncated 3’UTRs, as well as genes possessing RNA-seq reads but no NCBI annotated model, we sought to globally amend annotations to account for this. We first utilized cufflinks^80^ to generate transcript predictions using aligned RNA-seq reads with the options “--trim-3-dropoff-frac 0.25 --pre-mrna-fraction 0.3 --library-type fr-firststrand” on libraries from untreated subcaste RNA-seq data. Individual library-level transcript predictions were then merged using cuffmerge. These were then merged with NCBI annotations into a combined gff file following filtering out any transcripts with an FPKM < 10, and any cufflinks transcripts overlapping but distinct in exonic structure from NCBI transcripts were given the associated NCBI LOC ID# and included as alternative transcripts to the already-existing gene model. All cufflinks-identified gene models not represented by an NCBI gene model were then taken, and multi-exon predictions with an FPKM > 10 that did not overlap a known transcript on the opposite strand (cufflinks class_code “x” and “s”) were retained. For single-exon predictions, in order to further filter false positives, only single-exon models possessing a significant (1E-05) blast hit to fly were retained.

In order to create this adapted NCBI annotation, we first aligned all reads as above, but in the second round of alignment included the option “--outSAMattributes NH HI AS nM XS”. We then ran isoSCM v2.0.12 assemble with the options “-merge_radius 150 -s reverse_forward” on all alignments^81^. Following this we extracted only 3’ exons from the resulting draft annotations, which were then overlapped with the amended NCBI annotation. We then took the longest predicted 3’ exon/UTR that overlapped with the current NCBI version’s 3’ exon and extended this exon according to the isoSCM 3’ exon. Any predicted 3’ exon extension that overlapped an exon of another gene on the same strand was trimmed to 500 bp upstream of the other gene’s offending exon.

For all novel models (multi- and single-exon) transdecoder^82^ (v3.0.1) was used to predict coding sequence with the option “--single_best_orf”, and these were included when running reciprocal best blast on a combined annotation of NCBI, updated NCBI, and cufflinks novel genes.

### RNA-seq analysis

Read demultiplexing was performed using bcl2fastq2 (Illumina) with the options “--mask-short-adapter-reads 20 --minimum-trimmed-read-length 20 --no-lane-splitting --barcode-mismatches 0”. Reads were aligned to the *A. cephalotes* assembly using STAR^83^. STAR alignments were performed in two passes, with the first using the options “--outFilterType BySJout --outFilterMultimapNmax 20 --alignSJoverhangMin 7 --alignSJDBoverhangMin 1 --outFilterMismatchNmax 999 --outFilterMismatchNoverLmax 0.07 --alignIntronMin 20 --alignIntronMax 100000 --alignMatesGapMax 250000”, and the second using the options “--outFilterType BySJout --outFilterMultimapNmax 20 --alignSJoverhangMin 8 --alignSJDBoverhangMin 1 -- outFilterMismatchNmax 999 --outFilterMismatchNoverLmax 0.04 --alignIntronMin 20 --alignIntronMax 500000 --alignMatesGapMax 500000 --sjdbFileChrStartEnd [SJ_files]” where “[SJ_files]” corresponds to the splice junctions produced from all first pass runs.

Gene-level read counts were produced using featureCounts^84^ with the options “-O -M --fraction -s 2 –p”, against a custom adapted NCBI *A. cephalotes* annotation (see above). Resulting counts were rounded to integers prior to importing into R for DESeq2 analysis in order to account for non-integer count values associated with fractional counting of multi-mapping reads.

For all RNA-seq comparisons samples were taken from at least two distinct genetic backgrounds, and this was incorporated into all analyses to control for colony-of-origin effects. Differential gene expression tests were performed with DESeq2^85^. All pairwise comparisons were performed using the Wald negative binomial test (test = “Wald”) for determining differentially expressed genes, after blocking for colony background. Initially analyses were performed between behavioral groups (within and between subcastes) however upon observation of few transcriptomic differences between behavioral groups within a morphometric subcaste, samples were combined by morphometric group for following analyses. For morphometric subcaste-specific changes the specific subcaste was compared to all other samples in a given comparison. For comparisons between ‘outside’ and ‘nest’ subcastes, samples from the respective classes were tested against the other class.For tests of CCAP interaction effect (**Fig. 4E-F**), a likelihood ratio test approach was used in DESeq2, comparing the full model (genetic-background+caste+injection+[caste X injection]) to the reduced model lacking the interaction term (genetic-background+caste+injection), identifying genes that significantly differ in induction/repression response to CCAP injection depending upon subcaste-of-injection. Unless otherwise stated, an adjusted *P*-value cutoff of 0.05 was used to define differentially expressed genes.

### Statistics

Sample size and statistical tests are indicated in the figure legends. Unless otherwise noted, all statistical tests were two-sided. Boxplots were drawn using default parameters in R (center line, median; box limits, upper and lower quartiles; whiskers, 1.5x interquartile range). For testing significance of gene overlaps, Fisher’s exact tests were used after filtering out DEGs with “NA” for their adjusted p-value (indicating test failure due to low counts or variance) in either of the compared lists. For comparisons given in boxplots, non-parametric tests were used: Mann-Whitney U tests were used for all one-way comparisons, and Kruskal-Wallis test followed by Dunn’s correction for comparisons between 3 or more groups.

For comparisons with *H. saltator* and *C. floridanus* data, results from prior studies were linked to *A. cephalotes* genes via the reciprocal best blast hit method as outlined above^86,87^.

## Supplementary Table legends

**Table S1.** Results from gene ontology analysis of subcaste differential expression categories noted here. The 7^th^ column designates the specific subcaste or group tested and represents GO terms enriched among genes biased to that subcaste/group.

**Table S2.** Average transcripts per million, homology, and results of differential gene expression testing for *A. cephalotes* data here. For DEG calls empty cells indicate genes that lacked data (typically read counts) for testing, while “NS” indicates those with non-significant (padj ≥0.05) p-values.

**Table S3.** Proteomics results showing precursor charge, product charge, and fragments ions from identified proteins.

**Table S4.** Interacting protein with NPA following FLAG immunoprecipitation.

**Table S5.** Gene ontology results from experiments perturbing neuropeptides here. The 7^th^ column designates the specific perturbation and direction of tested DEGs and represents GO terms enriched among genes up- or down-regulated in the designated perturbation.

**Table S6.** Average transcripts per million, homology, and results from comparing nurse and forager *from H. glaber* as well as associated NPA application to cultured *H. glaber* astrocytes. For DEG calls empty cells indicate genes that lacked data (typically read counts) for testing, while “NS” indicates those with non-significant (padj ≥0.05) p-values.

**Table S7.** Results of all behavioral assays presented here.

**Video availability:** Example videos for every experiment and condition have been provided. We will upload all videos upon publication, but are currently attempting to find a suitable hosting site for public sharing of >500Gb of videos.

## Acknowledgements

We thank members of the Berger Lab (Charly Good and Khoa Tran) for help in editing the manuscript, Shirley Zeng, Christina Freeman, and Dave Schultz for providing help on peptide expression and purification. The 3D-printed behavioral chambers were courtesy of the University of Pennsylvania Library’s Biotech Common with design consultation. This work was supported by NIH training fellowships 5F31AG072777-03 (M.B.G.) and F32GM120933 (K.M.G), NIA R01s AG055570 (S.L.B.) and (AG047200). Live *A. cephalotes* colonies were collected in Trinidad in compliance with the laws of Trinidad and Tobago and imported to the USA in compliance with the conditions of the USDA APHIS Permit 526-23-94-85605A1.

## Author Contributions

K.M.G, M.F., M.B.G., and S.L.B. designed the experiments with input from M.S. and R.B. K.M.G., M.F. and M.B.G., collected the data. K.M.G and M.B.G. analyzed the data. T.G. performed the husbandry for the animals. D.X. created the primary neuronal cultures from leafcutter ant brains. J.B., L.K.P., and R.L. designed the LC-MS methods with oversight from B.A.G. A.B. and A.K. performed the naked mole-rat brain and cell experiments with guidance from A.S. and V.G. B.Z.K. set up the initial infrastructure for the leafcutter ant colonies. K.M.G., M.B.G., and S.L.B. wrote the manuscript with input from all co-authors. A.F. and M.W.M performed automated tracking analysis for CCAP peptide injection experiments.

---

**Figure S1.**
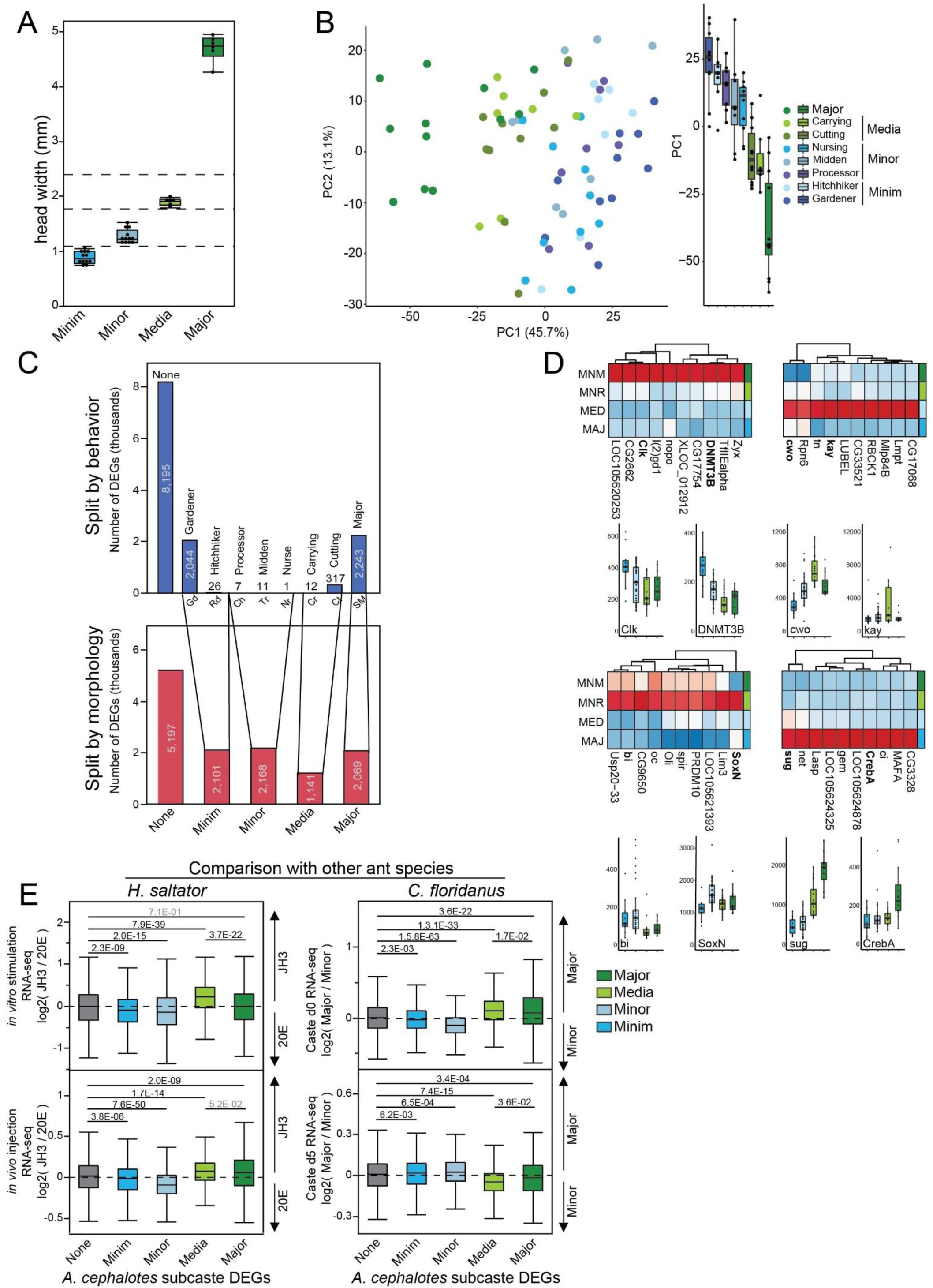
A) Measurements of maximum head width for each of the primary worker subcastes as determined here. B) PCA plot as in Fig 1B, but with all behavioral subcastes highlighted. C) Comparison of genes significantly elevated in a given behavioral group (top) as compared to the respective worker morphometric subcaste, and associated numbers of genes significantly elevated in (top numbers) the behavioral groups and (bottom) the four morphometric subcaste groups. Overall comparing behavioral groups resulted in far fewer condition-biased DEGs, while comparing morphometric castes (lower) showed strong transcriptomic differences. D) Top ten (by adjusted p-value from comparing each worker subcaste to all others) DNA binding domain containing genes significantly biased to each subcaste, with two representative genes’ expression shown in boxplots below (normalized counts). Upper left: Top ten Minim-biased TFs, upper right: top ten Media-biased TFs, lower left: top ten Minor biased TFs, lower right: top ten Major-biased TFs. E) Boxplots comparing genes biased to a given subcaste (relative to others) to log2-fold change of genes responding to JH3 or 20E in *H. saltator*^40^ either (top left) *in vitro* or (bottom left) *in vivo*, as well as (right) log2-fold change from comparing *C. floridanus* Major and Minor worker brain transcriptomes. This illustrates comparisons with general signatures of JH3/20E (left panels), as well as differences between a binary worker subcaste system as seen in *C. floridanus* (right), which we have previously associated with JH3 signaling^7–9^, overall suggesting persistent associations between worker caste and JH3/20E signaling in *A. cephalotes*. P-values generated by a Wilcoxon signed-rank test with Bonferroni correction following a significant Kruskal-Wallis test across all groups.

**Figure S2.**
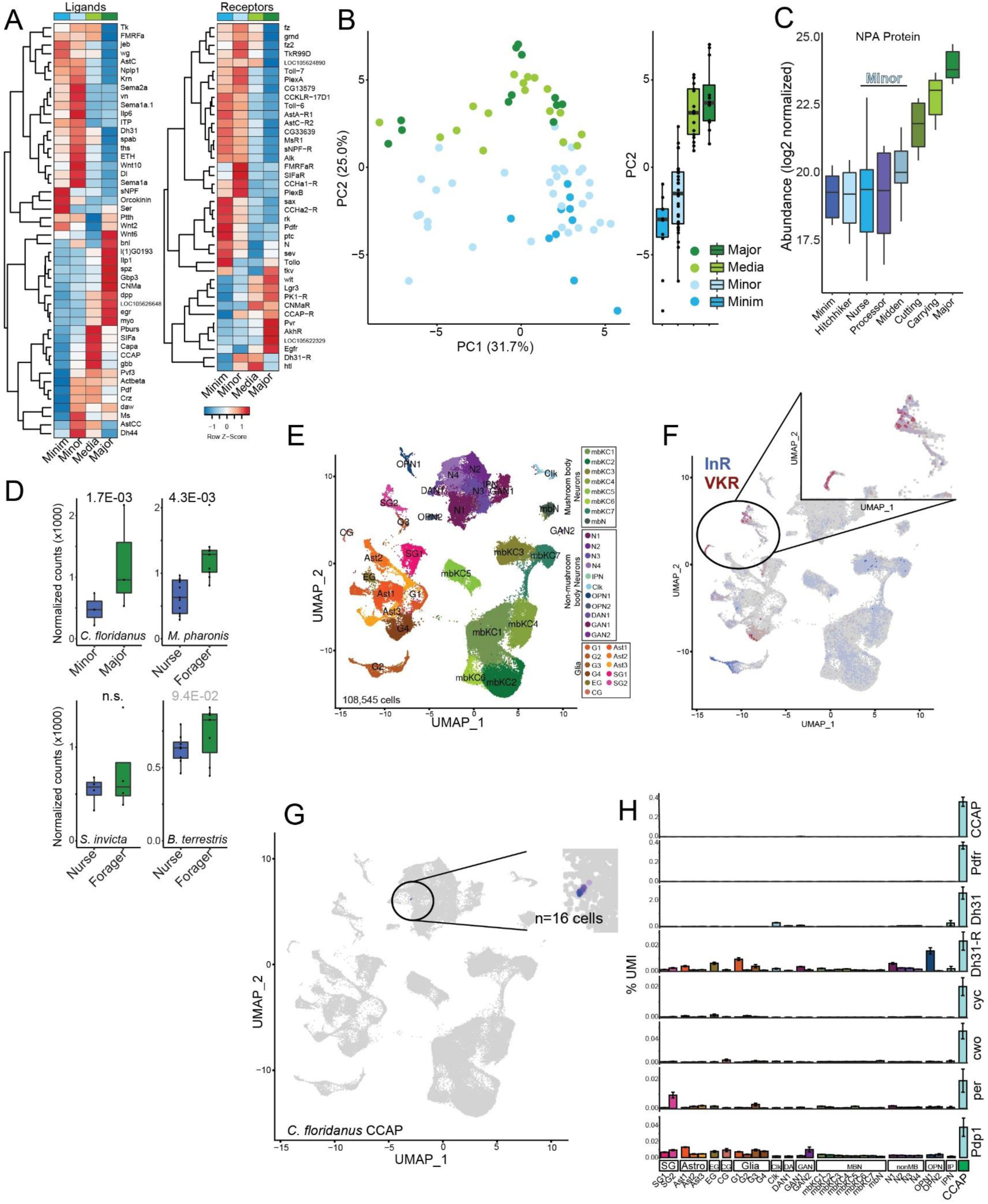
A) More inclusive list of ligands and receptors (as compared to Fig 2B), not limited to neuropeptides or neuropeptide-like genes. B) PCA analysis using only the genes presented in S2A (left) corresponding to peptide hormones, illustrating partial separation of subcaste based exclusively on this subset of genes. C) NPA Mass Spectrometry protein abundance (n=3). D) Normalized counts of NPA in four species showing significant or trending elevation in foraging subcastes in published data. E) UMAP plot from *C. floridanus* scRNA-seq^9^ illustrating cell types and cluster identities. F) UMAP showing InR and VKR co-expression within *C. floridanus* brain scRNA-seq data, illustrating co-expression particularly among cortex glia and perineurial surface glia (inset). G) UMAP plot showing cells expressing CCAP (expression > 4; 16 cells). H) Genes showing highest expression within cells expressing CCAP relative to all other clusters in *C. floridanus* d0 scRNA-seq clusters, illustrating circadian and neuropeptide co-expression with CCAP

**Figure S3.**
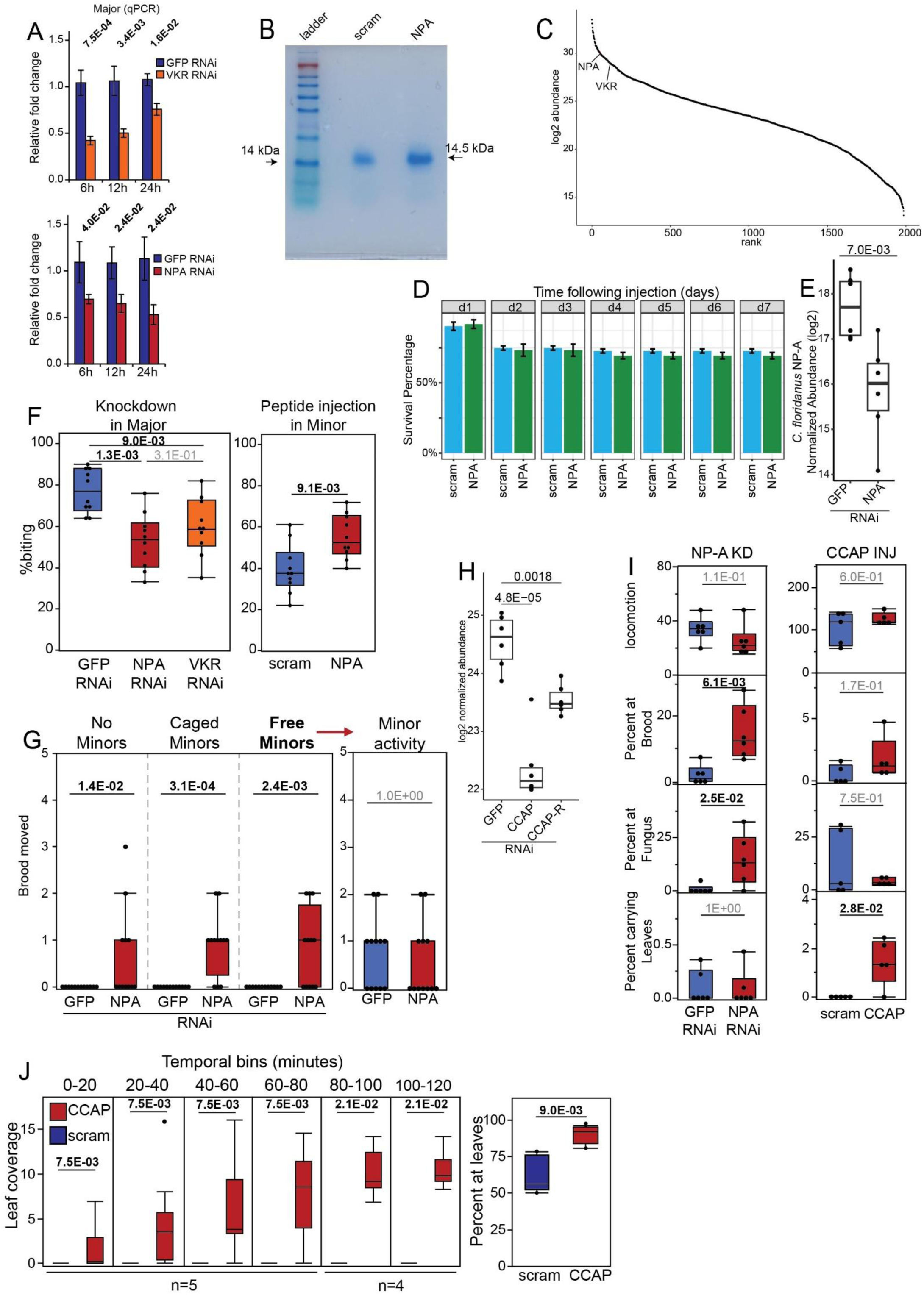
A) KD validation for NPA and VKR via RT-qPCR (n=6, each sample type). P-values generated by a Mann-Whitney U test. B) Coomassie gel to verify purification of scramble control peptide and recombinant NPA used here. C) IP-MS of NPA from Major brain showing VKR as top interactor and only receptor Immunoprecipitated with NPA (n=3). D) Mortality rates over 1-week for scramble control and NPA injection into brain (n=10). E) KD at protein level of NPA in *C. floridanus* (n=3, each sample type). P-values generated by a Mann-Whitney U test. F) Aggression assay (as in Fig 1B) performed for Majors following NPA or VKR KD as well as in Minor following NPA or scrambled control injection. NPA KD results in decreased aggression in Majors while NPA injection into Minors results in an increase in aggression. n=10 for all sample types. G) Social context NPA experiment illustrating that the presence of air-exposed caged Minor workers (typical caretaking subcaste) or Minors free within the assay does not impact Major acquisition of caretaking following NPA KD. Right: Minor worker brood movement from the ‘free Minor’ assay show no difference in caretaking despite increased Major brood interaction. n=12, each sample type. P-values generated by a Mann-Whitney U test. H) KD AT protein level of CCAP and CCAP-R (n=6, each sample type). P-values generated by a Mann-Whitney U test. I) Results from automated video tracking of NPA KD Majors (left, n=6 for all sample types) and CCAP peptide injected Majors (right, n=5 for all sample types). P-values generated by a Mann-Whitney U test. J) Binned automated assessment of Major leaf movement following CCAP peptide injection across 2 hours. Right: percent of time spent in leaf area overall for the same samples (n=5, each sample type). P-values generated by a Mann-Whitney U test of each pair.

**Figure S4.**
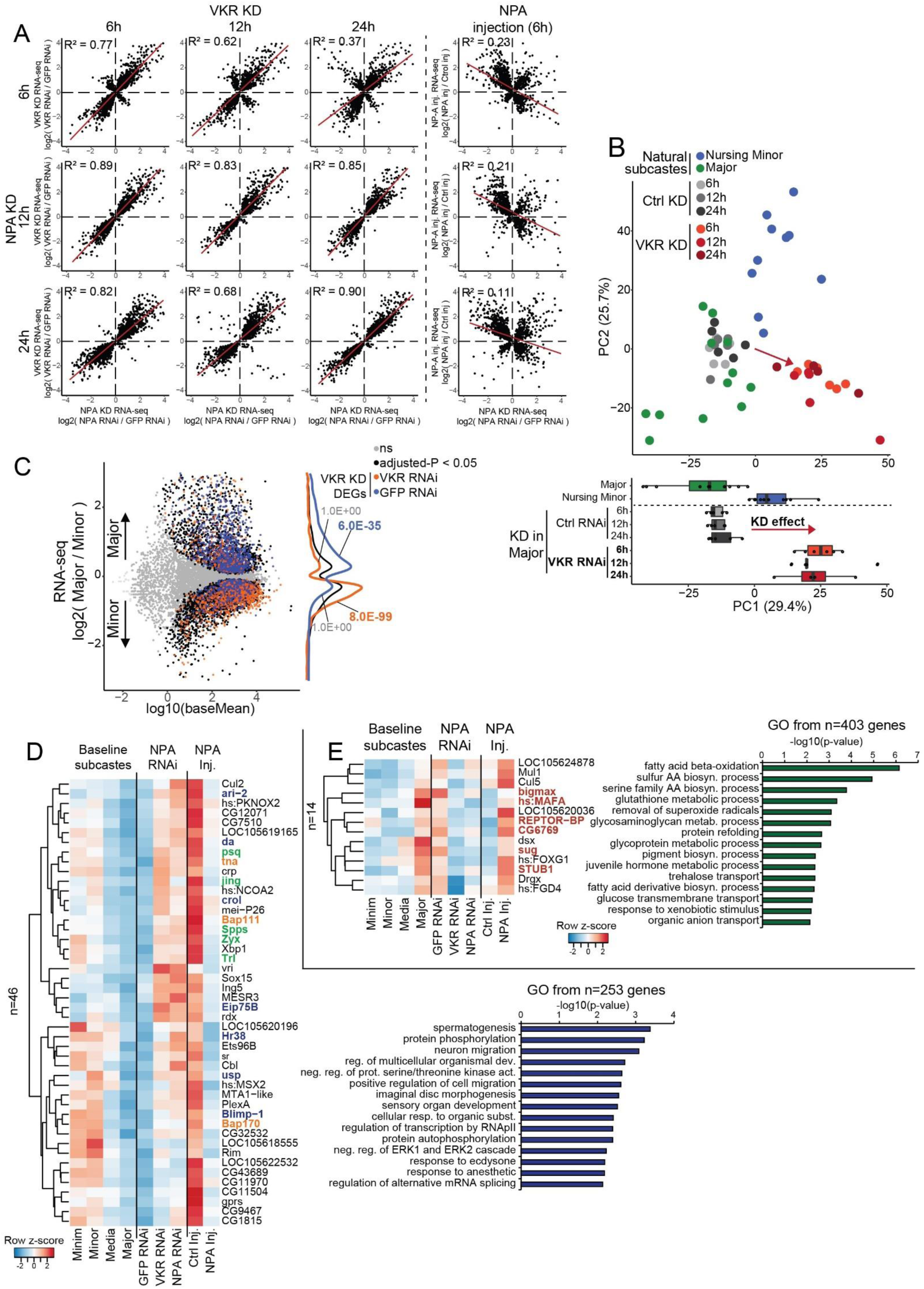
A) Correlation scatterplots for NPA KD timepoints (rows) as compared to VKR KD timepoints (columns). R2 represent pearsons correlations between log2 fold changes for each comparison. Overall this illustrates strong correspondence between NPA KD and VKR KD across timepoints. Rightmost column: NPA KD timepoints compared to 6h NPA injection, illustrating negative association between perturbations as would be expected if lowering NPA and increasing NPA lead to similar transcriptomic outcomes. B) PCA analysis of VKR (and GFP) KD samples as for Fig 4A. C) VKR KD DEGs overlapped with *A. cephalotes* Major vs Minor DEGs, illustrating that VKR KD in Major results in up-regulation of genes biased to Minor and down-regulation of genes biased to Major, as for NPA KD (Fig 4A). P-values represent the results of a fisher’s exact test comparing groups. D) Heatmap of all genes with DNA binding domain-like annotation showing elevated expression in Minor, significant up-regulation upon NPA and VKR KD, and significant down-regulation upon NPA injection. right: GO terms associated with *all* genes showing this pattern (n=253). Genes with key functions in fly colored: orange: brahma complex; green: Polycomb related; blue: Ecdysone response genes. E) Heatmap of all genes with DNA binding domain-like annotation showing elevated expression in Major workers, significant down-regulation upon NPA and VKR KD, and significant up-regulation upon NPA injection. right: GO terms associated with *all* genes showing this pattern (n=403). Red gene IDs represent genes related to regulation of energy homeostasis or metabolism.

**Figure S5.**
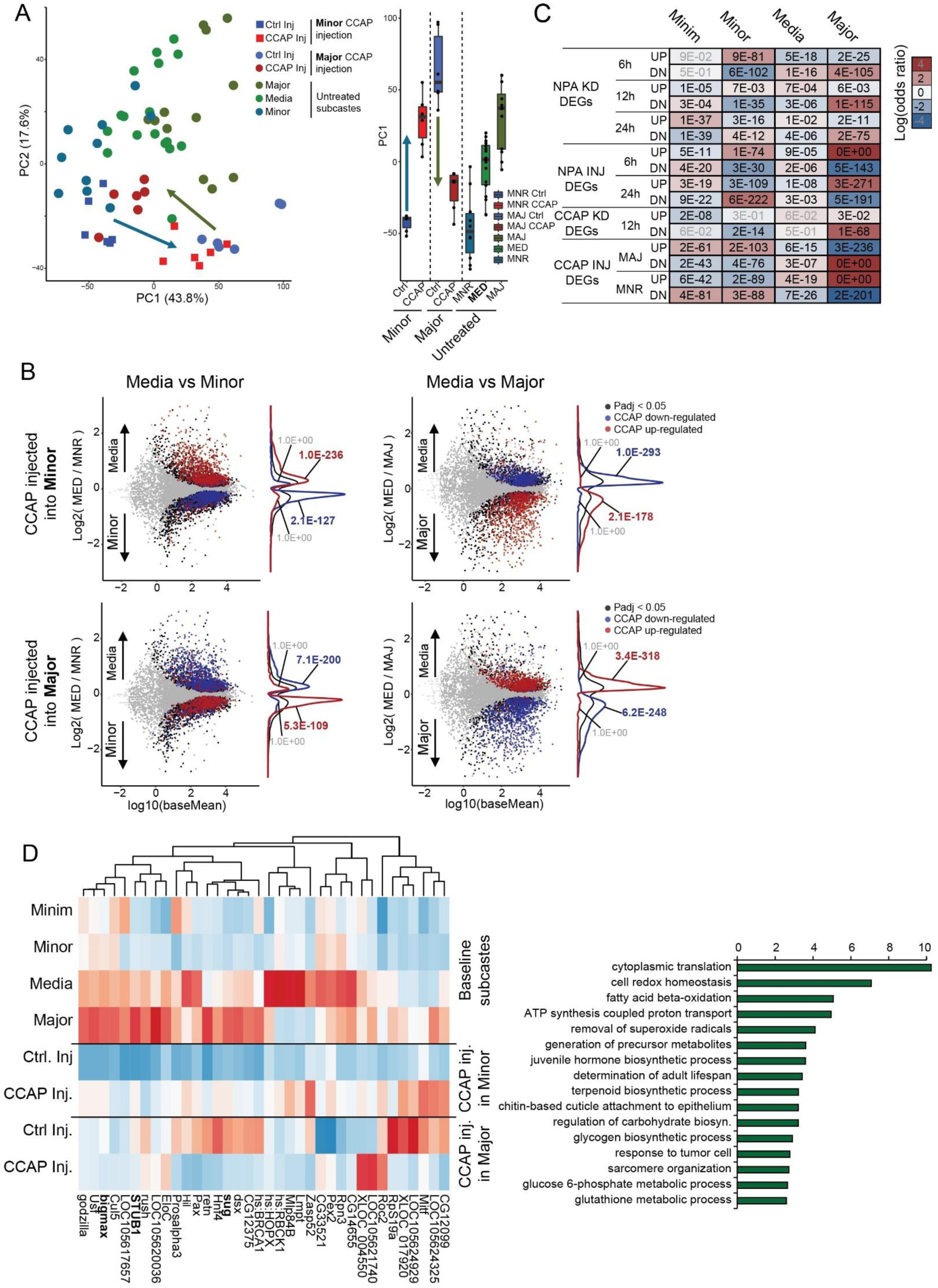
A) PCA of result from CCAP injection in Major and Minor as compared to respective controls as well as representative baseline subcastes, illustrating that in Minor CCAP injection (squares) results in a transcriptomic shift “Media-ward” and in Major the same but from opposite ends of PCA space. B) MA plots of baseline (left column) Media vs Minor and (right column) Media vs Major, overlain with (top row) Minor up- and down-regulated DEGs after CCAP injection, and (bottom roww) Major up- and down-regulated DEGs after CCAP injection into Major. Illustrating opposite effects that lead to up-regulation of genes typically elevated in Media, but only in the respective subcaste-of-injection. P-values represent the result of a fisher’s exact test comparing groups. C) Odds ratio (colors) and p-values from overlapping of perturbation DEGs with subcaste-biased DEGs for all comparisons in *A. cephalotes* here. P-values present the results of a fisher’s exact test comparing groups D) The same as for S4D but using genes featuring a DNA binding domain-like annotation and showing higher expression in Media relative to Minor, and upregulation upon CCAP injection into Minor. Right: gene ontology enrichment for all such genes (n=1018), regardless of DBD domain. Bolded gene IDs are those mentioned in the text.

**Figure S6.**
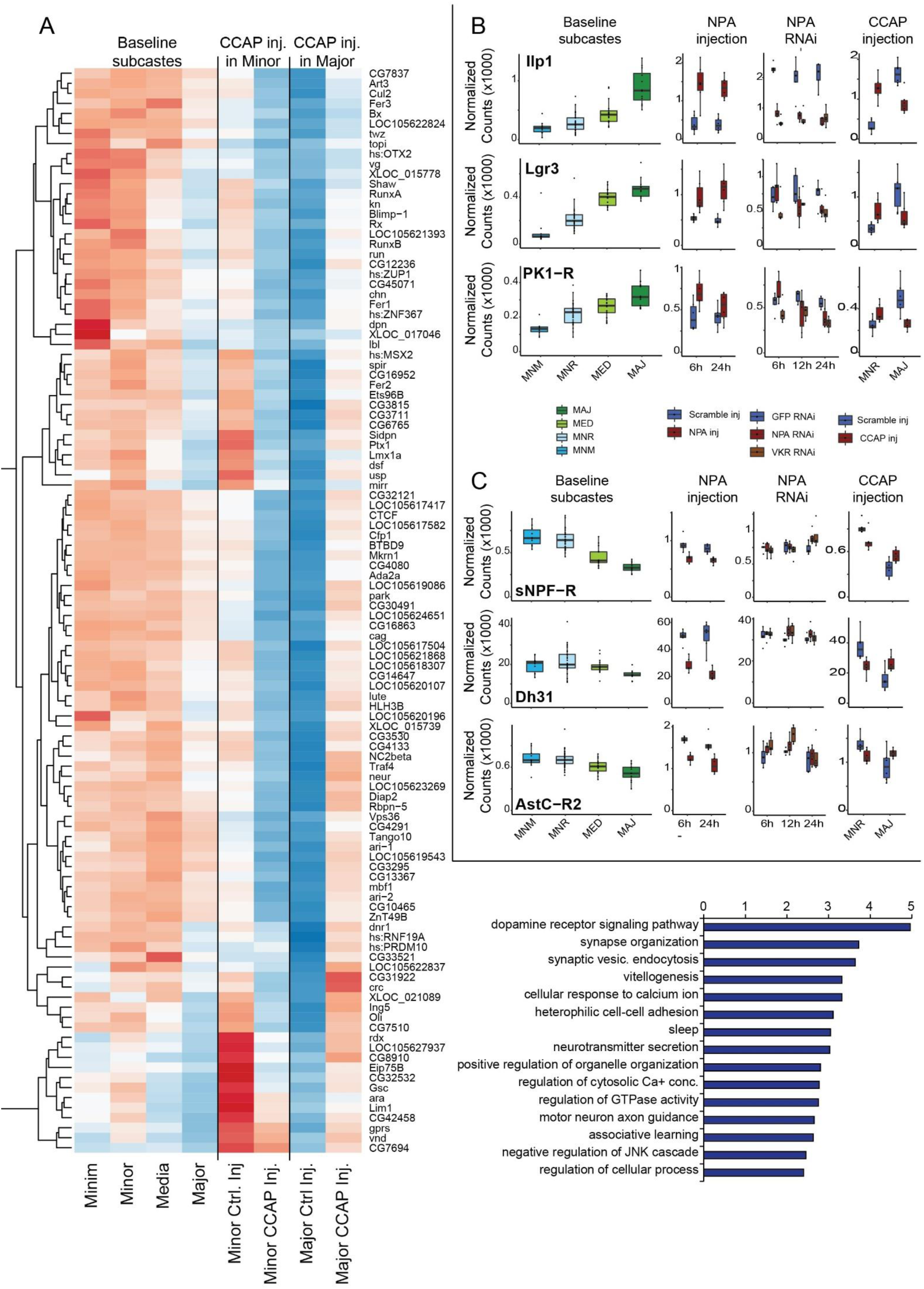
A) The same as for S4E but using genes featuring a DNA binding domain-like annotation and showing higher expression in Media relative to Major, and upregulation upon CCAP injection into Major. Right: gene ontology enrichment for all such genes (n=1,215), regardless of DNA binding domain-like annotation. B) Normalized count plots (in thousands) of neuropeptides or receptors showing bias to baseline Major and consistent regulation by NPA injection (middle left), NPA KD (middle right) and CCAP injection in reciprocal subcastes (rightmost). C) The same as S6B, but for those showing bias to baseline Minor/Minim.

**Figure S7.**
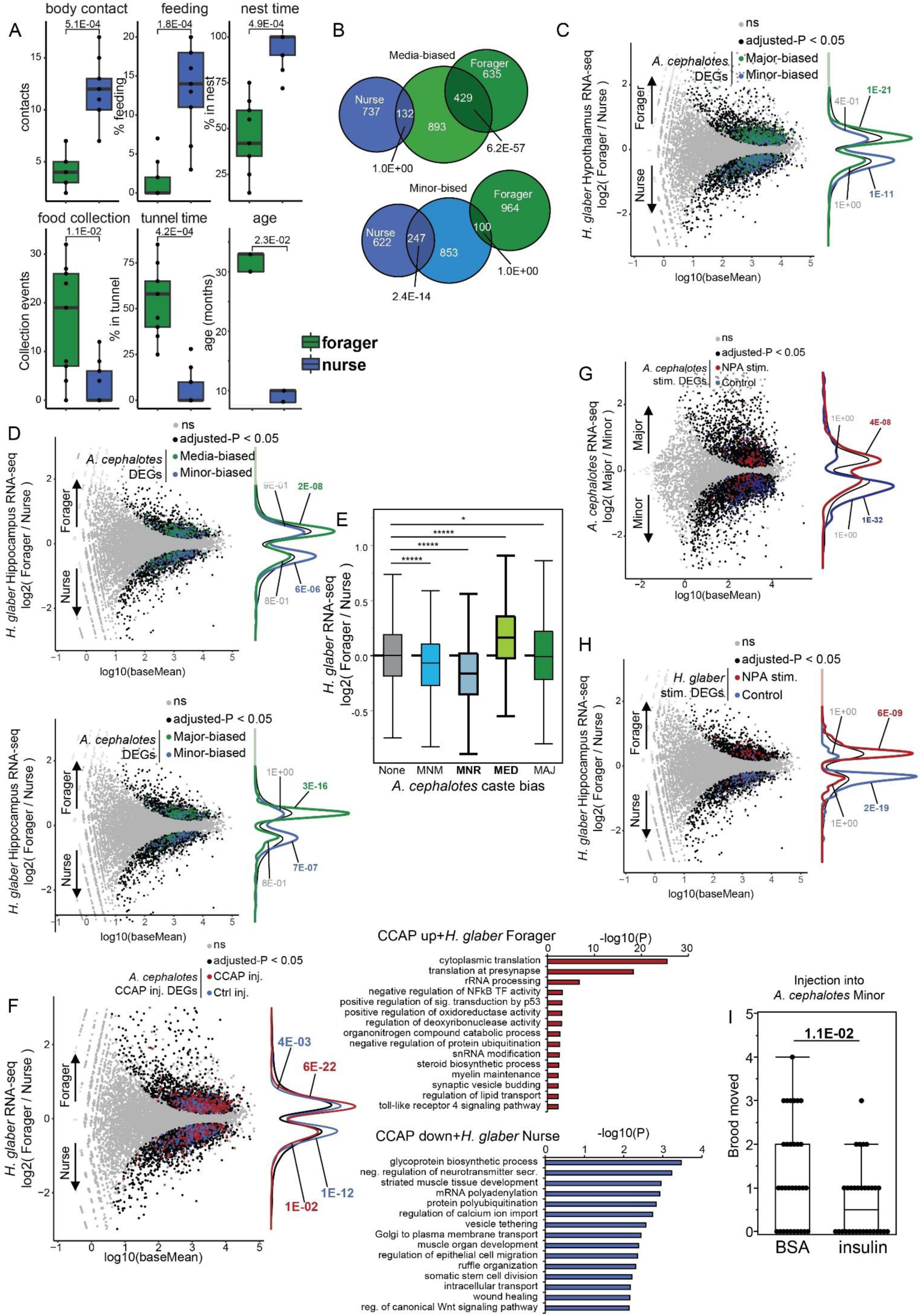
A) Metrics distinguishing *H. glaber* Nurse and Forager. Nurses were determined based upon body contacts with other non-breeders, time spent feeding pups (higher in nurses) and time residing within the local nest. Foragers were determined based upon age, food collection activity and time in non-proximal nest tunnels. P-values generated by a Mann-Whitney U test, n=9. Each group but age for which n=3. B) Venn diagram of the data shown in Fig 6B, showing numbers of genes. P-values represent the result of a fisher’s exact test comparing groups. C) Same comparison as for Fig 5B but overlain with **Major** vs Minor DEGs. P-values represent the result of a fisher’s exact test comparing groups. D) Same plots as figure 5B and S7B but for data from *H. glaber* Hippocampus, showing same trends. P-values represent the result of a fisher’s exact test comparing groups. E) Boxplots of *H. glaber* continuous subcaste bias (Forager / Nurse) as compared to major categories of subcaste genes in *A. cephalotes*, illustrating Minim and Minor genes show expression bias to *H. glaber* nurse, while Media show bias to *H. glaber* forager. P-values generated by a Mann-Whitney U test following a significant Kruskal Wallis test. F) Overlap between CCAP injection DEGs upon CCAP injection into *A. cephalotes* Minor with *H. glaber* hypothalamus differential expression. Right gene ontology terms associated with each class of gene overlap. P-values represent the result of a fisher’s exact test comparing groups. G) Overlap between *H. glaber* Forager vs Nurse gene expression with DEGs from *H. glaber* astrocyte NPA stimulation. P-values represent the result of a fisher’s exact test comparing groups. H) Overlap between *A. cephalotes* Major vs Minor gene expression and up- and down-regualted genes following NPA stimulation of *A. cephalotes* neuronal cultures. P-values represent the result of a fisher’s exact test comparing groups. I) Insulin injection into *A. cephalotes* Minor brain shows reduction of brood movement behavior. Assay performed as for Fig 3B, but using insulin (or BSA as control) injection (n=32, each sample type). P-values generated by a Mann-Whitney U test.

## Notes

### Competing Interest Statement

The authors have declared no competing interest.

